# The E2F4 transcriptional repressor is a key mechanistic regulator of colon cancer resistance to *irinotecan* (CPT-11)

**DOI:** 10.1101/2025.01.22.633435

**Authors:** Junichi Matsubara, Yong Fuga Li, Sanjay Koul, Junko Mukohyama, Luis E. Valencia Salazar, Taichi Isobe, Dalong Qian, Michael F. Clarke, Debashis Sahoo, Russ B. Altman, Piero Dalerba

**Affiliations:** Institute for Stem Cell Biology and Regenerative Medicine, Stanford University, Stanford, CA (USA); Department of Therapeutic Oncology, Graduate School of Medicine, Kyoto University, Kyoto (Japan); Department of Genetics, Stanford University, Stanford, CA (USA); Department of Bioengineering, Stanford University, Stanford, CA (USA); Illumina Inc., San Diego, CA (USA); Center for Discovery and Innovation (CDI), Hackensack Meridian Health (HMH), Nutley, NJ (USA); Department of Biological Sciences and Geology, Queensborough Community College (QCC), The City University of New York (CUNY), Bayside, NY (USA); Department of Pathology and Cell Biology, Columbia University, New York, NY (USA); Herbert Irving Comprehensive Cancer Center (HICCC), Columbia University, New York, NY (USA); Columbia Stem Cell Initiative (CSCI), Columbia University, New York, NY (USA); Department of Surgery, Institute of Medical Science, University of Tokyo, Tokyo (Japan); Department of Cell Biology, Albert Einstein College of Medicine, Bronx, NY (USA); Department of Comprehensive Oncology, Graduate School of Medicine, Kyushu University, Fukuoka (Japan); Department of Computer Science and Engineering, University of California San Diego (UCSD), San Diego, CA (USA); Department of Pediatrics, University of California San Diego (UCSD), San Diego, CA (USA), Department of Medicine (Division of Digestive and Liver Diseases), Columbia University, New York, NY (USA); Digestive and Liver Disease Research Center (DLDRC), Columbia University, New York, NY (USA); Department of Medical Sciences, Hackensack Meridian School of Medicine (HMSOM), Nutley, NJ (USA); Lombardi Comprehensive Cancer Center (LCCC), Georgetown University, Washington, DC (USA)

**Keywords:** colon cancer, cancer stem cells, cancer chemotherapy, drug resistance, irinotecan, E2F4

## Abstract

**Background.** *Colorectal carcinomas* (CRCs) are seldom eradicated by cytotoxic chemotherapy. Cancer cells with stem-like functional properties, often referred to as *“cancer stem cells”* (CSCs), display preferential resistance to several anti-tumor agents used in cancer chemotherapy, but the molecular mechanisms underpinning their selective survival remain only partially understood. **Methods.** In this study, we used *Transcription Factor Target Genes* (TFTG) enrichment analysis to identify transcriptional regulators (activators or repressors) that undergo preferential activation by chemotherapy in CRC cells with a *“bottom-of-the-crypt”* phenotype (EPCAM^+^/CD44^+^/CD166^+^; CSC-enriched) as compared to CRC cells with a *“top-of-the-crypt”* phenotype (EPCAM^+^/CD44^neg^/CD166^neg^; CSC-depleted). The two cell populations were purified in parallel by *fluorescence-activated cell sorting* (FACS) from a *patient-derived xenograft* (PDX) line representative of a moderately differentiated human CRC, following *in vivo* chemotherapy with *irinotecan* (CPT-11). The transcriptional regulators identified as differentially activated were tested for differential expression in normal vs. cancer tissues, and in cell populations enriched in stem/progenitor cell-types as compared to differentiated lineages (goblet cells, enterocytes) in the mouse colon epithelium. Finally, the top candidate was tested for mechanistic contribution to drug-resistance by selective down-regulation using *short-hairpin RNAs* (shRNAs). **Results.** Our analysis identified E2F4 and TFDP1, two core components of the DREAM transcriptional repression complex, as transcriptional modulators preferentially activated by *irinotecan* in EPCAM^+^/CD44^+^/CD166^+^ as compared to EPCAM^+^/CD44^neg^/CD166^neg^ cancer cells. The expression levels of both genes (*E2F4*, *TFDP1*) were found up-regulated in CRCs as compared to human normal colon tissues, and in a sub-population of mouse colon epithelial cells enriched in stem/progenitor elements (Epcam^+^/Cd44^+^/Cd66a^low^/Kit^neg^) as compared to other sub-populations enriched in either goblet cells (Epcam^+^/Cd44^+^/Cd66a^low^/Kit^+^) or enterocytes (Epcam^+^/Cd44^neg^/Cd66a^high^). Most importantly, E2F4 down-regulation using shRNAs dramatically enhanced the sensitivity of human CRCs to *in vivo* treatment with *irinotecan*, across three independent PDX models. **Conclusions.** Our data identified E2F4 and the DREAM repressor complex as critical regulators of human CRC resistance to *irinotecan*, and as candidate targets for the development of chemo-sensitizing agents.

## INTRODUCTION

*Colorectal carcinoma* (CRC) represents one of the most common forms of malignancy in the human population, accounting for approximately 9-10% of both cancer diagnoses and cancer-related deaths worldwide [1]. Approximately 15-20% of all CRC patients present at diagnosis with Stage IV disease (i.e., with overt distant-site metastases) and another 15-20% develop distant-site metastases as part of the disease’s natural progression, despite radical surgery to remove the primary tumor [2, 3]. To this date, cytotoxic chemotherapy, alone or in combination with anti-EGFR monoclonal antibodies, represents the standard of care for the first-line treatment of a majority of metastatic CRC patients, especially if affected by malignancies with a *microsatellite stable* (MSS) genotype [4, 5]. The efficacy of chemotherapy, however, is limited, resulting in 5-year *overall survival* (OS) rates of approximately 10% [6, 7]. In many CRCs, the malignant component of the tumor tissue is known to contain a variety of functionally distinct cell types, which recapitulate the various epithelial lineages found in normal colonic tissues, and which originate as a result of an epigenetic multi-lineage differentiation process, analogous to that sustained by normal stem cell populations as part of the homeostatic turnover of normal intestines [8, 9]. Included within these mixed populations of malignant, and clonally related, cells are also cellular elements that mirror the phenotype and transcriptional properties of normal stem/progenitor populations, and that appear capable of both self-renewal (i.e., serial propagation of tumor tissues in immune-deficient mice) and multi-lineage differentiation (i.e., systematic reconstitution, upon serial transplantation, of the cellular heterogeneity found in parent tissues) thus fulfilling the operational definition of *cancer stem cells* (CSCs) [8–11]. In many forms of cancer, CSC sub-populations have been shown to be preferentially resistant to a variety of anti-tumor agents [12–15]. However, the molecular mechanisms underlying such preferential resistance, especially resistance to cytotoxic drugs used in cancer chemotherapy, remain only partially understood [14, 15].

In previous studies, we observed that, in human CRCs, the cells that fulfill the functional definition of CSCs are restricted to a specific subset of the malignant cell population, defined by a surface marker phenotype (EpCAM^+^/CD44^+^/CD166^+^) that is characteristic of colon epithelial cells that are positioned at the *“bottom-of-the-crypt”*, where normal stem/progenitor populations are known to reside [8, 9]. In this study, we sorted EpCAM^+^/CD44^+^/CD166^+^ (CSC-enriched) and EpCAM^+^/CD44^neg^/CD166^neg^ (CSC-depleted) cancer cells from a *patient-derived xenograft* (PDX) line representative of a moderately differentiated human CRC, following *in vivo* treatment with either a placebo (saline solution) or *irinotecan* (CPT-11), an anti-tumor agent that stands at the backbone of systemic chemotherapy for metastatic CRCs [16, 17] and that is known to display only limited activity against human colon CSCs [12]. We then applied *Transcription Factor Target Gene* (TFTG) enrichment analysis, a bioinformatics technique designed to identify changes in the activity of transcription factors based on the coordinated change in expression levels of their target genes [18], in order to understand which transcriptional modulators are selectively (or preferentially) activated in CSC populations following chemotherapy. Our results indicate that, in human CRCs, exposure to *irinotecan* associates with preferential activation in CSCs of *E2F transcription factor 4* (E2F4) and *transcription factor dimerization partner 1* (TFDP1), two core components of the *dimerization partner* (DP)*, retinoblastoma-like* (RB-like)*, E2F and multi-vulva class B* (MuvB) *complex* (DREAM). Importantly, our experiments also show that E2F4 suppression with shRNA constructs sensitizes CRCs to *irinotecan*, dramatically enhancing the drug’s therapeutic activity.

## RESULTS

### Identification of the DREAM transcriptional repression complex as a mediator of the differential response observed in EpCAM^+^/CD44^+^/CD166^+^ as compared to EpCAM^+^/CD44^neg^/CD166^neg^ colon cancer cells following *in vivo* chemotherapy with *irinotecan* (CPT-11)

To investigate the effects of *irinotecan* on human colon CSCs, we engrafted into immune-deficient NOD/SCID/IL2R©^-/-^ (NSG) mice a PDX line representative of a moderately differentiated human CRC (PDX-COLON-8), and then treated tumor-bearing mice with *irinotecan*, administered according to a regimen designed to approximate those commonly used in patients with metastatic CRC (**Figure 1A-B**). We then used *fluorescence activated cell sorting* (FACS) to purify, side-by-side, two phenotypically distinct sub-populations of malignant cells known to co-exist in human CRC tissues (**Figure 1C**): 1) a population expressing surface markers characteristic of the *“bottom-of-the-crypt”* (stem/progenitor) compartment of colon epithelia (EpCAM^+^/CD44^+^/CD166^+^) and known to be enriched in CSC elements; and 2) a population expressing surface markers characteristic of the *“top-of-the-crypt”* (terminally differentiated) compartment of colon epithelia (EpCAM^+^/CD44^neg^/CD166^neg^) and known to be depleted in CSC elements [8, 9]. The two populations were first compared in terms of their rates of apoptotic death in response to drug treatment. The experiment confirmed that, indeed, the cytotoxic effects of *irinotecan* were largely restricted to the EpCAM^+^/CD44^neg^/CD166^neg^ (CSC-depleted, non-tumorigenic) cell population **(Supplementary Figure 1**). To gain insight into the molecular mechanisms responsible for such differential sensitivity, we used gene-expression microarrays (Affymetrix; platform: GPL15207) to profile the transcriptomes of the two populations, following treatment with either *irinotecan* or a negative control (saline solution). We then conducted an analysis by linear modeling to identify genes undergoing differential activation between the two populations, followed by a pathway-enrichment analysis designed to reveal whether such differentially modulated genes could be functionally clustered as contributing to the same biological processes [19, 20]. The analysis indicated that genes identified as differentially responsive to chemotherapy across the two populations were enriched for genes involved in cell-cycle regulation, DNA replication and *Aurora B kinase* (AURKB) signaling, all of which displayed an attenuated induction in the EpCAM^+^/CD44^+^/CD166^+^ (CSC-enriched) cell compartment **(Supplementary Table 1)**. We then applied TFTG enrichment analysis [18] to pinpoint individual transcription factors that might be responsible for such differential response in transcriptional induction. TFTG analysis identified as top candidates two transcriptional repressors: E2F4 and TFDP1 (**Figure 1D**). To better visualize this phenomenon, we conducted an aggregate analysis of the fold-change in expression levels of all analyzed genes, after sub-grouping into *“targets”* and *“non-targets”* of E2F4 and TFDP1 **(Supplementary Table 2)**. In accordance with our previous analysis, we observed that, although the average expression levels for target genes of E2F4 and TFPD1 were significantly induced by *in vivo* exposure to irinotecan in both EpCAM^+^/CD44^+^/CD166^+^ (CSC-enriched) and EpCAM^+^/CD44^neg^/CD166^neg^ (CSC-depleted) cells, such increase appeared substantially *“dampened”* in EpCAM^+^/CD44^+^/CD166^+^ (CSC-enriched) cells (**Figure 1E-F**). Because E2F4 and TFDP1 represent key mechanistic components of the same multi-protein complex, known as the DREAM transcriptional repression complex (**Figure 1G**), we interpreted our results as the manifestation of a preferential activation of such complex in EpCAM^+^/CD44^+^/CD166^+^ (CSC-enriched) cells.

**Figure 1.**
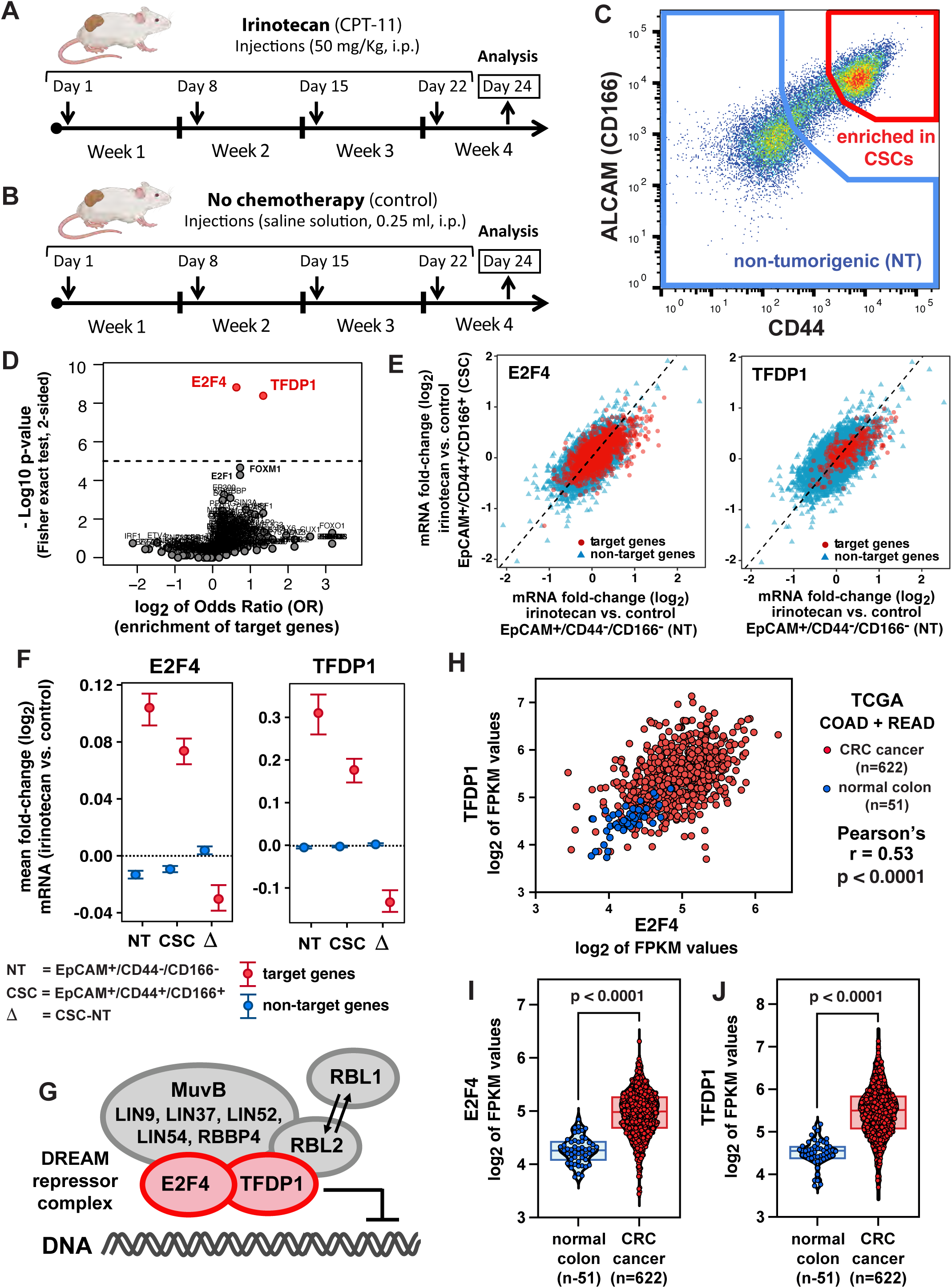
Identification of the E2F4 and TFDP1 transcriptional repressors, two core members of the DREAM complex, as candidate mechanistic regulators of the differential response to irinotecan of colon cancer cells with a “*cancer stem cell*” (CSC) phenotype. **(A-B)** Schematic representation of the treatment regimen used to evaluate the *in vivo* anti-tumor activity of irinotecan (CPT-11). Irinotecan was administered to tumor-bearing mice based on a dosing schedule (50 mg/kg, once weekly x 4 weeks) designed to approximate the pharmacokinetics of the drug’s active metabolite (SN-38) in human colon cancer patients. Irinotecan’s activity was compared to that of a negative control, consisting of the drug’s vehicle alone (saline solution). Downward arrows: drug injections. Upward arrows: tumor analysis. Images created using BioRender.com. **(C)** Scatter-plot showing the expression profile of CD44 and ALCAM (CD166) in a PDX-COLON-8 tumor, as evaluated by flow cytometry. The scatter-plot is gated on live human epithelial cells (DAPI^neg^, Mouse-lineage^neg^, EpCAM^+^) and displays the co-existence of two major sub-populations: 1) a population expressing high levels of CD44 and ALCAM (CD44^+^/CD166^+^) and known to be enriched in cells with a *“cancer stem cell”* (CSC) phenotype (red gate); and 2) a population expressing low levels of CD44 and ALCAM (CD44^neg^/CD166^neg^) and known to consist in *non-tumorigenic* (NT) cells (blue gate). **(D)** Volcano plot reporting the results of the *transcription factor target gene* (TFTG) analysis, in which each *transcription factor* (TF) is plotted based on the magnitude of its differential activation in EpCAM^+^/CD44^neg^/CD166^neg^ as compared to EpCAM^+^/CD44^+^/CD166^+^ cells following *in vivo* exposure to irinotecan (x-axis) and the statistical significance of such differential activation (y-axis). Dotted line: p=0.00001 (cutoff for statistical significance after Bonferroni correction). The volcano plot identifies two transcriptional repressors (E2F4, TFDP1) among the most differentially activated TFs (red dots). **(E)** Scatter-plots showing the fold-change in expression level following *in vivo* exposure to irinotecan of all measurable genes, as measured in EpCAM^+^/CD44^neg^/CD166^neg^ cells (x-axis) as compared to EpCAM^+^/CD44^+^/CD166^+^ cells (y-axis), and after stratification of genes in *targets* (red circles) and *non-targets* (blue triangles) of E2F4 and TFDP1. Following *in vivo* exposure to irinotecan, genes suppressed by E2F4 and TFDP1 display a higher induction in EpCAM^+^/CD44^neg^/CD166^neg^ (NT) as compared to EpCAM^+^/CD44^+^/CD166^+^ (CSC) cells, as revealed by their visual enrichment below the line of equivalence (x=y; dotted line). **(F)** Average fold-change in expression level following *in vivo* exposure to irinotecan, and average difference in such fold-change (Δ), for genes classified as targets and non-targets of E2F4 and TFDP1, in EpCAM^+^/CD44^neg^/CD166^neg^ cells (NT) as compared to EpCAM^+^/CD44^+^/CD166^+^ cells (CSC). The difference in the up-regulation of E2F4 and TFDP1 target genes is statistically significant (Welch’s t-test, 2-sided). Error bars: mean +/-95% *confidence interval* (CI). **(G)** Schematic representation of the DREAM repressor complex, listing its major structural components. The complex can include, alternatively, the RBL2 (p130) or RBL1 (p107) proteins. **(H)** Scatter-plot displaying the distribution of *E2F4* and *TFDP1* mRNA expression levels in normal colorectal tissues (n=51) and primary CRCs (n=622) included in the *Colon Adenocarcinoma* (COAD) and *Rectal Adenocarcinoma* (READ) datasets of *The Cancer Genome Atlas* (TCGA) repository. E2F4 and TFDP1 expression levels are positively correlated to each other (r=0.56; p <0.001). **(I-J)** Violin-plots comparing the distribution of *E2F4* and *TFDP1* expression levels between normal colorectal tissues (n=51) and primary CRCs (n=622) from TCGA’s COAD and READ datasets. Both *E2F4* and *TFDP1* are overexpressed in tumor as compared to normal tissues (Mann-Whitney U-test, two-sided).

### Increased expression of *E2F4* and *TFDP1* mRNAs in *“bottom-of-the-crypt”* (stem/progenitor) compartments of normal colon epithelia

To understand whether an enhancement of the capability to activate the DREAM complex was to be considered a general feature of malignant tissues, we first tested whether the expression levels of *E2F4* and *TFDP1* mRNAs were higher in CRCs as compared to normal colonic tissues. To answer this question, we combined the *Colon Adenocarcinoma* (COAD) and *Rectal Adenocarcinoma* (READ) datasets of *The Cancer Genome Atlas* (TCGA), which include a large collection of RNA-seq experiments performed on paired specimens of primary CRCs (n=622) and normal colonic epithelia from adjacent mucosal tissues (n=51). The analysis showed that the mRNA expression levels for *E2F4* and *TFDP1* were positively correlated to each other (**Figure 1H**) and higher in malignant as compared to normal tissues (**Figure 1I-J**), in agreement with the notion that, in human CRCs, the fraction of cells with a EpCAM^+^/CD44^+^/CD166^+^ phenotype tends to increase as compared to normal colon epithelia [8, 9]. To understand whether increased expression of genes encoding for core elements of the DREAM complex was indeed characteristic of a subset of colon epithelial cells with a stem/progenitor phenotype, we decided to quantify the mRNA expression levels for *E2f4* and *Tfdp1* across different subtypes of epithelial cells in the normal mouse colon, where the molecular identity and surface marker phenotype of stem/progenitor cells are better understood [21–24]. We used *RNA-sequencing* (RNA-seq) to study the transcriptional profile of three distinct sub-populations of normal mouse colon epithelial cells, purified by FACS from primary tissues: 1) a *“bottom-of-the-crypt”* population enriched in both *Lgr5^+^ columnar basal cells* (CBCs) as well as other epithelial cell types with intestinal stem/progenitor properties, such as *Fgfbp1^+^*cells (Epcam^+^/Cd44^+^/Cd66a^low^/Kit^neg^); 2) a *“bottom-of-the-crypt”* population enriched in goblet cells (Epcam^+^/Cd44^+^/Cd66a^low^/Kit^+^); and 3) a *“top-of-the-crypt”* population enriched in terminally differentiated enterocytes (Epcam^+^/Cd44^neg^/Cd66a^high^) (**Figure 2A, Supplementary Figure 2**). The analysis revealed that, indeed, expression of both *E2f4* and *Tfdp1* was higher in the population enriched in stem/progenitor cells (Epcam^+^/Cd44^+^/Cd66a^low^/Kit^neg^) as compared to those enriched in goblet cells or terminally differentiated enterocytes (**Figure 2B-C**).

**Figure 2.**
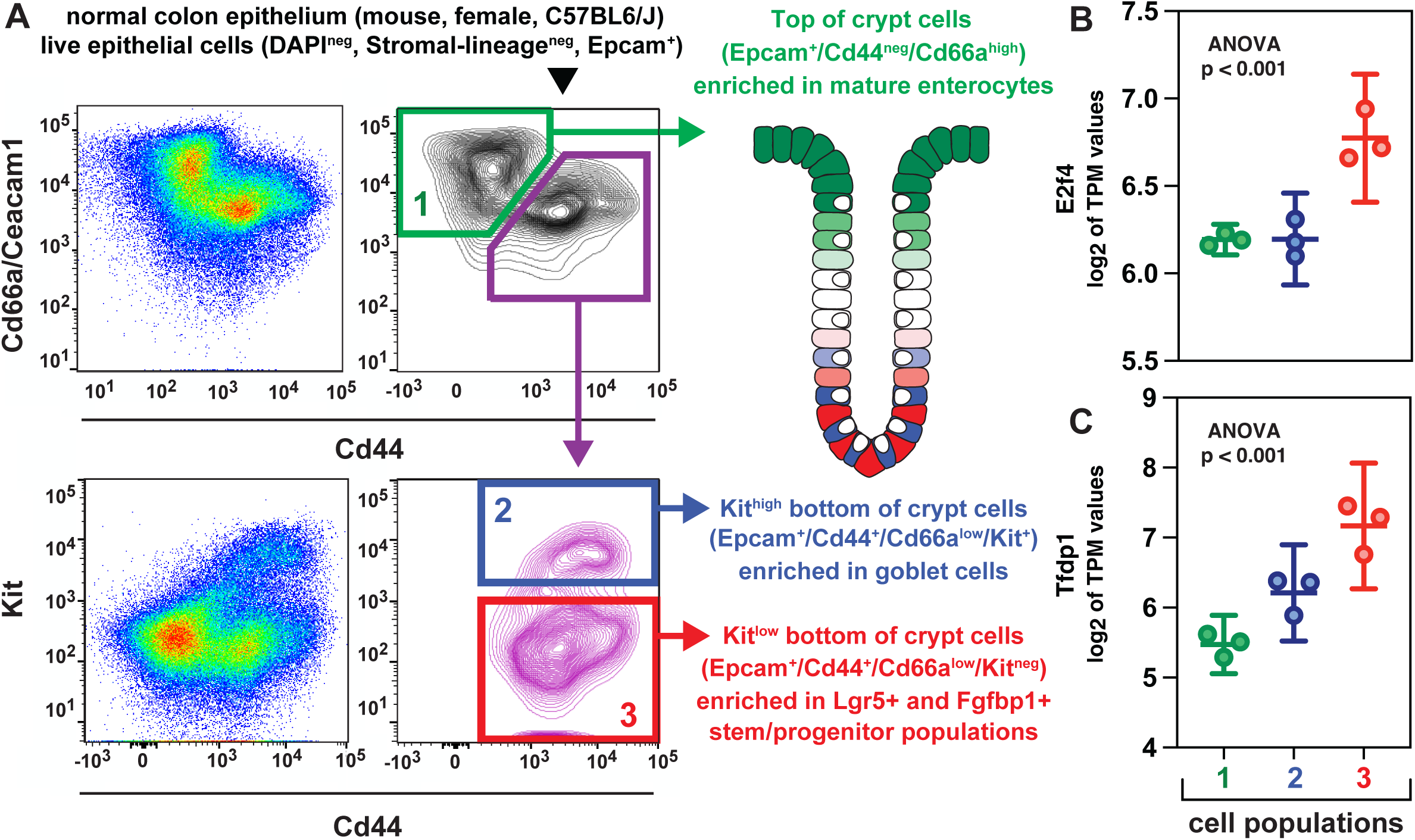
Differential expression of *E2f4* and *Tfdp1* across distinct epithelial cell types within the mouse colonic epithelium. **(A)** Schematic representation of the FACS strategy used to purify three distinct phenotypic subsets of epithelial cells from the colonic mucosa of adult C57BL6/J female mice. The scatter-plots are gated on live cells (DAPI^neg^) lacking expression of surface markers restricted to stromal cell-types (Stromal-lineage^neg^) but expressing a surface marker characteristic of epithelial cells (Epcam^+^). The three sub-types of colon epithelial cells sorted by FACS correspond to: 1) Epcam^+^/Cd44^neg^/Cd66a^high^ cells, enriched in terminally differentiated enterocytes located at the top of colonic crypts; 2) Epcam^+^/Cd44^+^/Cd66a^low^/Kit^+^ cells, enriched in goblet cells located at the bottom of colonic crypts; and 3) Epcam^+^/Cd44^+^/Cd66a^low^/Kit^neg^ cells, enriched in both Lgr5^+^ and Fgfbp1^+^ cell-types located at the bottom of colonic crypts and known to include cells with stem/progenitor properties. A complete description of the sorting strategy, and a listing of the surface markers used to exclude stromal cell-types, is provided in **Supplementary Figure 2. (B-C)** Analysis by RNA-seq of *E2f4* and *Tfdp1* mRNA expression levels across the three different phenotypic subsets of mouse normal colon epithelial cells purified by FACS as described in Panel A. Both *E2f4* and *Tfdp1* are overexpressed in Epcam^+^/Cd44^+^/Cd66a^low^/Kit^neg^ cells (enriched in Lgr5^+^ and Fgfbp1^+^ stem/progenitor cell-types) as compared to Epcam^+^/Cd44^+^/Cd66a^low^/Kit^+^ cells (enriched in goblet cells) and Epcam^+^/Cd44^neg^/Cd66a^high^ cells (enriched in terminally differentiated enterocytes), as evaluated by a one-way Welch ANOVA. Error bars: mean +/-95% CI.

### E2F4 is a mechanistic regulator of colon cancer sensitivity to *irinotecan*

To understand whether, in human CRCs, the DREAM repression complex played a mechanistic role in causing the increased resistance of EpCAM^+^/CD44^+^/CD166^+^ (CSC-enriched) cancer cells to *in vivo* treatment with *irinotecan* (CPT-11), we decided to test whether E2F4 knock-down by *short hairpin RNA* (shRNA) constructs would translate in increased CRC sensitivity to the drug, across a variety of PDX lines **(Supplementary Table 3)**. We obtained from *The RNAi Consortium* (TRC) of the *Broad Institute* two shRNA constructs designed to target E2F4 (E2F4-shRNA[#1]: TRCN0000013808; E2F4-shRNA[#2]: TRCN0000013809) **(Supplementary Table 4)** [25], and then cloned them into lentivirus vectors designed to express them in a constitutive fashion, in tandem with a *copepod-derived green fluorescent protein* (copGFP) reporter that enabled purification of lentivirus-infected cells by FACS. When constitutively expressed in CRC cells using lentivirus vectors, both shRNA constructs proved capable of down-regulating E2F4 expression, at both the mRNA and protein level **(Supplementary Figure 3**). Because E2F4 and the DREAM complex are known to modulate the transcription of several genes involved in cell-cycle regulation [26, 27], and because previous studies reported E2F4 to play, alternatively, a positive or a negative role on cell proliferation depending on specific experimental systems [28], we decided to test whether E2F4 down-regulation would translate in a functional perturbation of the proliferative capacity of EpCAM^+^/CD44^+^/CD166^+^ (CSC-enriched) cancer cells at baseline (i.e., in the absence of chemotherapy). We purified EpCAM^+^/CD44^+^/CD166^+^ (CSC-enriched) cancer cells from two PDX lines engineered to express E2F4-shRNAs (PDX-COLON-8, PDX-COLON-60) and then tested them for quantitative reductions in their capacity to initiate the formation of either copGFP^+^ *three-dimensional* (3D) organoids *in vitro* or solid tumors *in vivo*, as compared to isogenic controls engineered to express an empty lentivirus vector **(Supplementary Figure 4**). In our experimental system, E2F4 knock-down did not translate into a reduction in the frequency of organoid-initiating cells (*in vitro*) or the frequency of tumor-initiating cells (*in vivo*) as evaluated by an *extreme limiting dilution assay* (ELDA). This observation was consistent with the general consensus on the biology and function of E2F4 and the DREAM complex, which envisions them a transcriptional repressors of the cell-cycle machinery [28] that become actively recruited to either maintain cellular quiescence (in the G_0_-phase of the cell-cycle) or induce cellular arrest (in the G_2_-phase of the cell cycle) [28–31], based on a variety of environmental stimuli and functional contexts.

Finally, we tested whether E2F4 knock-down, despite not directly impacting the tumorigenic capacity of CRC cells, would nonetheless render them more vulnerable to *in vivo* chemotherapy with *irinotecan*, administered according to the same dosing regimen that we used to investigate drug-induced transcriptional perturbations and designed to approximate the dosing regimens used in human patients (**Figure 1A-B**). In a first experiment, we tested whether E2F4 knock-down would translate into an increased induction of apoptosis in malignant PDX tissues, either at baseline or following treatment with *irinotecan* (**Figure 3**). Our results showed that, in the absence of treatment, tumors engineered to express E2F4-specific shRNA constructs did not associate with a measurable increase in the activation of pro-apoptotic effector pathways (cleaved Caspase-3^+^) within the malignant component (copGFP^+^) of cancerous tissues, as compared to control tumors (i.e., tumors infected with an empty lentivirus vector). In the presence of chemotherapy with irinotecan, on the other hand, tumors engineered to express E2F4-specific shRNA constructs displayed a substantial increase in the percentage of cancer cells undergoing apoptosis, substantially higher than that observed in controls receiving the same treatment (mean percentage of apoptotic cancer cells within tissues infected with an empty lentivirus vector vs. E2F4-shRNA[#1] vs. E2F4-shRNA[#2]: 3.8% vs. 12.9% vs. 14.8%; Dunn’s test for pairwise comparisons: p<0.0001; 2-way ANOVA test for interaction between E2F4 knock-down and chemotherapy: p<0.0001). Finally, we tested whether E2F4 knock-down would translate into increased tumor sensitivity to *irinotecan* (**Figure 4**). In this case we compared the *in vivo* anti-tumor activity of irinotecan against three independent PDX lines, engineered to expressed either an empty lentivirus backbone (baseline control) or the E2F4-shRNA[#2] construct (which displayed a slightly superior activity as compared to construct E2F4-shRNA[#1], in terms of both suppression of E2F4 expression and *in vivo* CRC sensitization to irinotecan-induced apoptosis). The results showed that, in the absence of treatment, CRCs engineered to express the E2F4-shRNA construct displayed similar (if not higher) growth kinetics as compared to isogenic controls. In the presence of chemotherapy with irinotecan, on the other hand, CRCs engineered to express the E2F4-shRNA construct displayed a substantial increase in drug sensitivity, as revealed by the systematic observation of profound reductions in average tumor volumes and growth rates, superior in magnitude to those observed in control tumors (i.e., statistically significant in 2-way and 3-way ANOVA tests for interaction between E2F4 knock-down and chemotherapy) and often associated with prolonged tumor shrinkage (which is usually not observed when using irinotecan alone).

**Figure 3.**
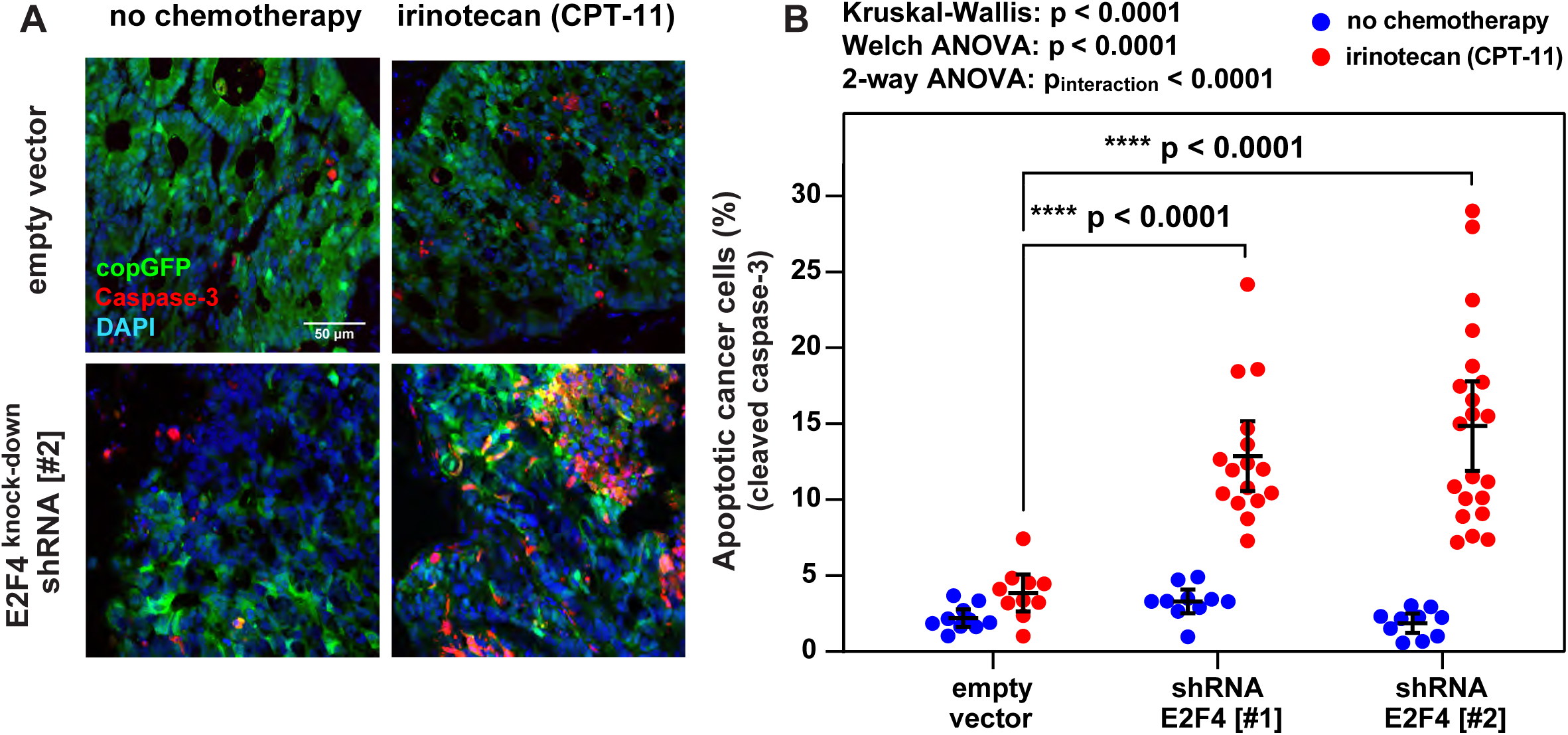
Effects of E2F4 knock-down on the *in vivo* sensitivity of colon cancer to irinotecan, as evaluated by the induction of apoptosis in tumor tissues. **(A)** Analysis by *immunofluorescence* (IF) of cleaved Caspase-3 expression (red) in PDX-COLON-8 tumor tissues engineered to express a green, fluorescent reporter (copGFP), either alone (empty vector) or in tandem with a *short hairpin* RNA (shRNA) construct designed to knock-down E2F4 (#2), following *in vivo* treatment with either irinotecan (50 mg/kg, once weekly x 4 weeks) or a negative control (saline solution). Cell nuclei are counterstained with DAPI (blue). Scale bar: 50 μm. **(B)** Quantification of the percentage of apoptotic cancer cells observed in PDX-COLON-8 tumor tissues following E2F4 knock-down with two distinct shRNA constructs (#1, #2) and *in vivo* treatment with irinotecan. The percentage of apoptotic cancer cells was computed by dividing the number of apoptotic cancer cells (copGFP^+^, Caspase-3^+^) by the total number of cancer cells (copGFP^+^) within the same microscopic field. For each experimental condition, a minimum of 10 independent microscopic fields were examined, representative of a minimum of 4 independent tumors. E2F4 knock-down resulted in a statistically significant increase in the percentage of apoptotic cancer cells following *in vivo* treatment with irinotecan, as evaluated using a non-parametric test (Kruskal-Wallis H-test + Dunn’s test for pairwise comparisons), a parametric test (Welch ANOVA + Dunnet’s T3 test for pairwise comparisons) and a test for interaction between E2F4 knock-down and *in vivo* treatment with irinotecan (two-way ANOVA + Dunnett’s test for pairwise comparisons). Error bars: mean +/-95% CI.

**Figure 4.**
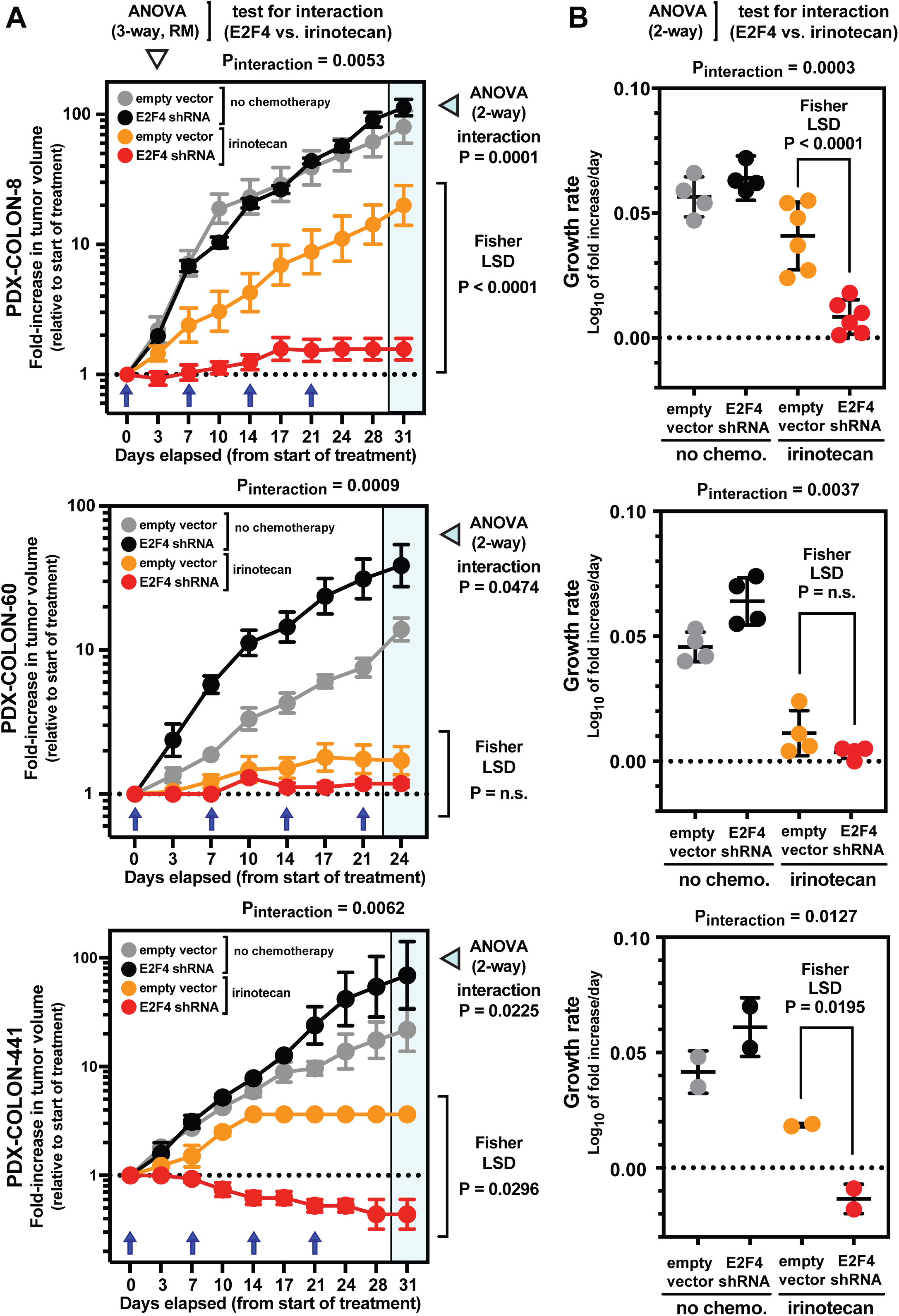
Effects of E2F4 knock-down on the *in vivo* sensitivity of colon cancer to irinotecan, as evaluated by the induction of objective tumor responses and/or reduction of growth rates. **(A-F)** Comparison of average growth kinetics, expressed as either fold-increases in average tumor volumes (A, C, E) or growth rates (B, D, F), between tumors infected with a lentivirus vector encoding for an shRNA construct designed to knock-down E2F4 expression [#2] and tumors infected with an empty lentivirus vector (negative control), following *in vivo* treatment with either irinotecan (50 mg/kg, once weekly x 4 weeks) or a negative control (saline solution). The experiment was repeated on three distinct *patient-derived xenograft* (PDX) lines established from three distinct *colorectal carcinomas* (CRCs): PDX-COLON-8 (A-B), PDX-COLON-60 (C-D), PDX-COLON-441 (E-F). In all three PDX models, E2F4 knock-down resulted in a substantial enhancement of irinotecan’s anti-tumor activity, as indicated by profound suppression of growth rates and, in one model, induction of objective tumor responses. The role of E2F4 knock-down in enhancing tumor sensitivity to irinotecan was evaluated for statistical significance using a test for interaction (E2F4 shRNA vs. irinotecan) within the framework of a 3-way ANOVA for *repeated measures* (RM) over time (time x lentivirus vector x treatment) and a 2-way ANOVA (lentivirus vector x treatment) on both fold-increases in tumor volumes at the last time-point and growth rates, followed by a Fisher *least significant difference* (LSD) test for the pre-determined comparison (empty vector + irinotecan vs. E2F4 shRNA + irinotecan). Growth rates were calculated assuming exponential growth kinetics. Error bars for fold-increases in tumor volume: mean +/-*standard error of the mean* (SEM). Blue arrows: days at which treatment was administered (chemotherapy vs. saline solution). Error bars for growth rates: mean +/-*standard deviation* (SD).

## DISCUSSION

In this study, we identified E2F4, a key mechanistic component of the DREAM transcriptional repression complex, as an important regulator of CRC sensitivity to *irinotecan* (CPT-11), a derivative of *camptothecin* (CPT) and a key anti-tumor drug used as standard-of-care for metastatic CRC [16, 17]. In our experimental system, suppression of E2F4 expression translated into a dramatic enhancement of the *in vivo* anti-tumor activity of *irinotecan*, across multiple pre-clinical models of human CRC.

Although the biological reason behind this association remains unknown, this finding appears coherent with a substantial wealth of knowledge on the mechanism of action of *irinotecan* and the biological functions of E2F4. Upon systemic administration, *irinotecan* is converted into SN-38 by *carboxyl-esterases* (CE) found in the liver [32]. Like CPT and many of its analogues, SN-38 is a potent poison of the human *topoisomerase-1* (TOP1) enzyme [33, 34], an endonuclease responsible for both introducing *single-strand breaks* (SSBs) in DNA molecules and then repairing them by re-ligation [35]. Among the key functions of TOP1 is to prevent the progressive *“supercoiling”* of *double-stranded* DNA (dsDNA) during replication, by allowing it to *“unwind”* by *“swiveling”* around one of its two strands (i.e., the non-cleaved strand) down-stream of replicative forks [35]. When poisoned by CPT, TOP1 molecules become unable to re-ligate the SSBs that they previously introduced in the DNA, and thus become *“paralyzed”* at cleavage sites, in the form of *DNA-protein cross-links* (DPC) in which the catalytic domain of TOP1 is linked by a covalent bond to the 3’ end of the SSB [35–37]. In proliferating cells that are actively progressing through the S-phase of the cell cycle, TOP1 inhibition by CPT results in the accumulation of widespread *double-strand DNA breaks* (DSBs) because of *“collisions”* between advancing replicative forks and stalled TOP1-DPCs at cleavage sites, ultimately resulting in cell death [36, 37]. Importantly, however, because both TOP1-induced SSBs and TOP1 inhibition by CPT are rapidly reversible [38, 39], a transient inhibition of cell proliferation shortly before exposure to CPT is able to protect cancer cells from the drug’s cytotoxic effects [40, 41], especially when CPT is administered for a relatively short period of time and at lower concentrations, in a manner that mirrors the pharmacokinetics of SN-38 exposure *in vivo* [42, 43].

E2F4 and the DREAM transcriptional repression complex, on the other hand, are known to play a central role in cell-cycle regulation [28], as they contribute to suppress, in an aggregate and coordinated fashion, the expression of a large and diverse set of genes that are mechanistically required for cell-cycle progression [26, 27], thus acting as *“enforcers”* of both cellular quiescence (e.g., maintenance of cells in the G_0_-phase of the cell-cycle) [28] and cellular arrest (e.g., inhibition of cell-cycle progression beyond the G_2_-phase following exposure to DNA damaging agents) [29, 30]. A possible mechanistic interpretation of our observations, therefore, could envision E2F4 and the DREAM complex as cell-cycle *“guardians”*, tasked with protecting the genetic integrity and ensuring the preferential survival of adult stem-cell populations, either by limiting their exposure to the mutagenic effects of DNA-damaging agents (e.g., by enforcing a baseline quiescence in the G_0_-phase of the cell cycle) [28] or by limiting their apoptotic death following exposure to DNA-damaging agents during the S-phase of an active cell-cycle (e.g., by delaying S-phase progression and/or inducing G_2_-phase arrest, thus allowing time for DNA-repair) [29, 30]. In the specific case of CRCs treated *in vivo* with *irinotecan*, and thus transiently exposed to low doses of SN-38 (10-100 nM), a preferential activation of E2F4 and the DREAM complex in CSC populations with a *“bottom-of-the-crypt”* phenotype (EPCAM^+^/CD44^+^/CD166^+^) would be expected to allow for their preferential survival, as it would delay their progression through the S-phase, allowing time for both the spontaneous reversion of TOP1-induced SSBs before the collision of TOP1-DPCs with replicative forks and the robust induction of enzymatic systems for the repair of DSBs. This interpretation is coherent with our observations on the distribution of *E2F4* and *TFDP1* mRNA expression levels in mouse and human intestinal tissues, which revealed that: 1) within the mouse colonic epithelium, *E2f4* and *Tfdp1* mRNA expression levels are highest in the subset of cells with a *“bottom-of-the-crypt”* non-secretory phenotype (Epcam^+^/Cd44^+^/Cd66a^low^/Kit^neg^), which are known to be enriched in stem/progenitor populations [21], including both Lgr5^+^ [22] and Fgfbp1^+^ cell-types [23, 24]; and 2) *E2F4* and *TFDP1* mRNA expression levels are higher in human CRCs as compared to normal colonic epithelia, in agreement with the notion that, in human CRCs, the malignant epithelium is enriched in cells with a *“bottom-of-the-crypt”*, stem/progenitor phenotype (EPCAM^+^/CD44^+^/CD166^+^) when compared to normal colon epithelia from the same patients [8, 9]. This interpretation is also coherent with multiple lines of evidence form the published literature, including: 1) the phenotype of *E2f4*^-/-^ mice, which display reduced body size and abnormal development of the intestinal epithelium when compared to wild-type littermates [44]; 2) studies on the role of E2F4 in modulating the sensitivity of prostate cancer cells to radiotherapy, showing extensive DNA fragmentation in the presence of E2F4 down-regulation by *small interfering RNAs* (siRNAs) [29]; and 3) studies on the functional effects of spontaneous E2F4 somatic mutations found in CRCs with *microsatellite instability* (MSI), displaying an extension of the protein’s half-life without apparent compromise of its modulatory activity on transcription [45].

In terms of translational applications, the results of our study indicate that, in human CRCs, E2F4 and the DREAM complex could represent an important therapeutic target to enhance the clinical efficacy of state-of-the-art therapeutic regimens. Based on our observations, pharmacological manipulations capable of inhibiting the signaling pathways that mediate assembly and/or maintenance of the DREAM complex in its repressor configuration would be predicted to phenocopy the effects caused by E2F4 downregulation, and thus strongly sensitize metastatic CRCs to *irinotecan*. Among the possible candidates for this type of application are the inhibitors of *dual specificity tyrosine phosphorylation regulated kinase 1A* (DYRK1A), an enzyme that contributes to maintain the repressor configuration of the DREAM complex by phosphorylating one of its component proteins (LIN52) [31]. Indeed, recent studies have reported that genetic knock-out of the *DYRK1A* gene sensitizes CRC cells to *topotecan* [46], an anti-tumor drug similar to *irinotecan* in chemical structure and mechanism of action, commonly used to treat ovarian cancer. Small-molecule pharmacological inhibitors of DYRK1A, such as harmine, are actively being developed for the treatment of diabetes, because they promote expansion of insulin-secreting pancreatic ®-cells [47] and appear well-tolerated in both animal models and human volunteers [47–49]. Harmine itself, for example, is found at relatively high concentrations (1-2 mg/ml) in psychoactive herbal brews such as *ayahuasca,* which is used for spiritual and medicinal purposes in many cultures indigenous to South America [49]. Our study, therefore further advocates for the development of pre-clinical studies aimed at testing DYRK1A inhibitors as potential chemo-sensitizers, to enhance the anti-tumor activity of *irinotecan* against human models of CRC. Finally, as mentioned above, *irinotecan* belongs to a class of chemotherapeutic agents (i.e., CPT-derivatives) that include several compounds already approved for clinical use, either formulated as individual drugs (e.g., *irinotecan*, *topotecan*) [16, 17, 50] or as part of *antibody-drug conjugate* (ADC) molecules (e.g., *deruxtecan*, *govitecan*) [51]. Because camptothecin-derivatives are widely used in medical oncology to treat different tumor types (e.g., colon cancer, ovarian cancer, breast cancer), and because E2F4 and the DREAM complex represent fundamental components of the cell-cycle machinery, expressed ubiquitously across many organs and tissues, our findings are likely to be applicable across a wide spectrum of clinical settings, and cold be leveraged to improve the therapeutic activity of chemotherapy for a relatively large and diverse population of cancer patients.

## METHODS

### Ethics statement

All animal experiments were conducted in compliance with research protocols pre-approved by the *Administrative Panel on Laboratory Animal Care* (APLAC) of *Stanford University* (protocol number: 10868) and *Institutional Animal Care and Use Committee* (IACUC) of *Columbia University* (protocol numbers: AC-AAAW1466, AC-AABM9553). All newly established *patient-derived xenograft* (PDX) lines were originated from malignant tissues specimens obtained from colon cancer patients that provided informed consent, in compliance with a research protocol pre-approved by the *Institutional Review Board* (IRB) of *Stanford University* (protocol number: 4344) [8, 9]. All researchers performing procedures that involved the use of human specimens were pre-approved by the IRBs of *Stanford University* (protocol number: 4344) and *Columbia University* (protocol number: AAAT0960).

### Establishment and propagation of *patient-derived xenograft* (PDX) lines

Human PDX lines were established by direct *sub-cutaneous* (s.c.) implantation into immune-deficient NOD/SCID/IL2Rγ^-/-^ (NSG) mice (Charles River Laboratories: strain code #614; The Jackson Laboratory: stock #005557) of solid tissue fragments collected from primary tissue specimens of human *colorectal carcinomas* (CRCs), following previously published protocols and procedures [8, 9]. A list of the PDX lines used in this study is provided as part of **Supplementary Table 3**, alongside relevant elements of the clinical and pathological information relative to corresponding patients (e.g., sex, age, disease stage, site of primary origin, pathological grade). Briefly, primary human CRC tissue specimens were minced with scissors into small (2 mm^3^) fragments and implanted *sub-cutaneously* (s.c.) using a 10-gauge Trochar needle, through a small incision on the animal’s right dorsal flank. Recipient mice (6-8 weeks old, female) were briefly anesthetized by isoflurane inhalation (AErrane®, Baxter Healthcare Corporation) using a standard vaporizer (5% for induction, 2% for maintenance). Skin incisions were closed using *VetBond* (3M). Once established, solid tumor xenografts were serially propagated using the same technique.

### *In vivo* chemotherapy experiments

Immediately following implantation with human colon cancer PDX lines, immune-deficient NSG mice were randomly assigned into treatment (*irinotecan*) or control (*saline solution*) groups and then treated as soon as *sub-cutaneous* (s.c.) tumors became palpable. *Irinotecan* (CPT-11) was purchased as a clinical-grade formulation in liquid solution (20 mg/ml; Camptosar®, Pfizer), then diluted 1:5 to the desired concentration (4 mg/ml) using saline solution (0.9% NaCl) and finally administered *intra-peritoneally* (i.p.) to tumor-bearing mice, at a dose of 50 mg/kg of mouse body weight, once weekly x 4 weeks (**Figure 1A**). This treatment regimen was designed to fulfill three criteria: a) to mirror a dosing schedule commonly used for the administration of *irinotecan* in human patients undergoing chemotherapy for metastatic CRC [16, 17]; b) to administer a dose of *irinotecan* that, among those recommended for pre-clinical studies in mice [52, 53], would be conservatively positioned at the lower end, in order to enable a better recapitulation of the plasma concentrations observed in human patients for SN-38, the active metabolite of *irinotecan* [32, 42]; and c) to use an administration route that would minimize experimental variability and distress to animals, without compromising the drug’s bio-distribution and pharmacokinetics [54]. Animals belonging to control groups received i.p. injections of saline solution (0.25 ml), according to a schedule identical to that used for irinotecan (**Figure 1B**). Tumor size was measured by caliper twice weekly, and tumor volumes estimated using the formula recommended by Tomayko and Reynolds [55]:

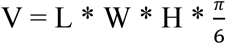

In the above formula: V = *tumor volume*; L = *tumor length*; W = *tumor width*; H = *tumor height*. Mice were euthanized either at the conclusion of the 4-week treatment course (day 24) or when tumors reached 20 mm in maximum diameter (our ethical endpoint for early euthanasia), whichever occurred first. In experiments designed to test the role of E2F4 knock-down in the modulation of cancer resistance to *irinotecan*, tumors were generated by s.c. injection in NSG mice of human CRC cells infected with lentivirus constructs encoding for *short-hairpin RNAs* (shRNAs) designed to knock-down E2F4 expression. In all cases, before injection, human CRC cells were purified by *fluorescence activated cell sorting* (FACS) based on their differential expression of a *copepod green fluorescent protein* (copGFP) reporter, encoded by the lentivirus vectors in tandem with the shRNAs, in order to ensure that all transplanted cells would consist of genetically engineered cells, irrespective of infection efficiency. Lentivirus-infected (copGFP^+^) human CRC cells were injected s.c. at a dose known to result in robust engraftment (i.e., 10,000 cells/mouse). The role of E2F4 knock-down in enhancing tumor sensitivity to irinotecan was evaluated for statistical significance using a test for interaction (E2F4 shRNA vs. irinotecan) within the framework of a 3-way ANOVA for *repeated measures* (RM) over time (time x lentivirus vector x treatment) and of a 2-way ANOVA (lentivirus vector x treatment), performed on both fold-increases in tumor volumes at each experiment’s last time-point as well as growth rates, followed by a Fisher *least significant difference* (LSD) test for the pre-determined comparison (empty vector + irinotecan vs. E2F4 shRNA + irinotecan). Growth rates were calculated assuming exponential growth kinetics [56–58].

### Dissociation of solid tissues into single-cell suspensions

The procedures for the dissociation of solid tissues into single-cell suspensions were based on previously published protocols, and were tailored to the specific type of tissue being processed, either human CRC xenografts [8, 9] or normal mouse colonic tissues [21, 59]. Briefly, in the case of human CRC xenografts [8, 9], s.c. tumor lesions were harvested from NSG mice, minced into small fragments (2 mm^3^) using scissors, then rinsed with *Hank’s balanced salt solution* (HBSS), finely chopped with a razor blade into minute fragments (0.2-0.5 mm^3^), resuspended in RPMI-1640 medium supplemented with L-alanine-L-glutamine (2 mM; GlutaMAX, Invitrogen), penicillin (120 μg/ml), streptomycin (100 μg/ml), amphotericin-B (0.25 μg/ml; Invitrogen), HEPES (20 mM; Sigma-Aldrich), sodium pyruvate (1 mM; Lonza), DNase-I (100 units/ml; Worthington) and Collagenase-III (200 units/ml; Worthington) and finally incubated for 2 hours at 37 °C, to obtain enzymatic disaggregation into a single-cell suspension. Cell suspensions were gently dissociated by mechanical pipetting, then serially filtered through a sterile gauze, a 70-μm nylon mesh and a 40-μm nylon mesh, in order to remove undigested tissue fragments and large cell clumps. Red blood cells were removed by osmotic lysis, achieved by incubating the cell suspension in *ammonium chloride potassium* (ACK) hypotonic buffer (NH_4_Cl, 150 mM; KHCO_3_, 1 mM) for 5 min on ice (4°C). In the case of normal mouse colonic tissues [21, 59], analyses were conducted on the distal segment of the colon and rectum (i.e., the segment of the colon distal to the caecum, which was excluded). Colonic tissues were harvested from adult mice (6-8 weeks old, female) of the C57BL/6J strain (The Jackson Laboratory; stock #000664). Colonic tissues were opened longitudinally, rinsed with cold *Dulbecco’s Phosphate Buffered Saline* (DPBS) solution to remove fecal material, and finally incubated in DPBS supplemented with EDTA (5 mM), for 45 min on ice (4°C). During incubation with EDTA (5 mM), colonic tissues were gently stirred, to facilitate the detachment of epithelial crypts. Epithelial crypts floating in liquid suspension were spun-down by centrifugation, resuspended in RPMI-1640 medium supplemented with L-alanine-L-glutamine (2 mM; GlutaMAX, Invitrogen), penicillin (120 μg/ml), streptomycin (100 μg/ml), amphotericin-B (0.25 μg/ml; Invitrogen), HEPES (20 mM; Sigma-Aldrich), sodium pyruvate (1 mM; Lonza), DNase-I (100 units/ml; Worthington) and Collagenase-III (200 units/ml; Worthington), and finally incubated at 37 °C for 30 minutes, to obtain enzymatic disaggregation into a single-cell suspension. Dissociated cells were re-suspended by gentle pipetting and serially filtered, first with sterile gauze and subsequently with 70-μm and 40-μm nylon meshes, in order to remove large cell clumps. Contaminating red blood cells were removed by osmotic lysis with ACK hypotonic buffer (NH_4_Cl, 150 mM; KHCO_3_, 1 mM) for 5 min on ice (4°C).

### Flow cytometry and *fluorescence-activated cell sorting* (FACS)

To minimize loss of cell viability, all FACS experiments were conducted on fresh single-cell suspensions, prepared shortly before their analysis by flow cytometry, following established protocols and procedures [8, 9, 21, 59]. In all experiments, antibody staining was performed in HBSS supplemented with 2% *heat-inactivated calf serum* (HICS), EDTA (5 mM), penicillin (120 μg/ml), streptomycin (100 μg/ml), amphotericin-B (0.25 μg/ml), HEPES (20 mM) and sodium pyruvate (1 mM). The protocols for the differential visualization of distinct sub-types of colon epithelial cells were tailored to the specific tissue being analyzed, either human CRC xenografts [8, 9] or normal mouse colonic tissues [21, 59]. Briefly, in the case of human CRC xenografts [8, 9], cells were first incubated with human IgGs (0.6% w/v; Gammagard Liquid, Baxter) for 20 min on ice (4°C), at a concentration of 3-5 × 10^5^ cells/100 μl, in order to “block” human *Fc receptors* (FcRs) and therefore minimize unspecific binding of antibodies on human cells. Cells were subsequently washed and stained with antibodies at dilutions determined by appropriate titration experiments. The antibodies used to visualize cancer cells with *bottom-of-the-crypt* (EpCAM^+^, CD44^+^, CD166^+^) and *top-of-the-crypt* (EpCAM^+^, CD44^neg^, CD166^neg^) phenotypes included: anti-human-EpCAM-AlexaFluor488 (clone 9C4; BioLegend), anti-human CD44-PE-Cy7 (clone G44-26; BD Biosciences) and anti-human-CD166-PE (clone 105902; R&D Systems). A visual illustration of the gating strategy implemented to sub-group human epithelial cells (EpCAM^+^) into *bottom-of-the-crypt* (CD44^+^, CD166^+^) and *top-of-the-crypt* (CD44^neg^, CD166^neg^) phenotypic subsets is provided in **Figure 1C**. Mouse cells (Mouse-lineage^+^) were excluded by staining with anti-mouse H-2K^d^-PacificBlue (clone SF1-1.1; BioLegend). In experiments aimed at the detection of early apoptotic cells, cells were also stained with Annexin-V-APC (BioLegend), in conjunction with the reagents and instructions provided by the *FITC-Annexin-V Apoptosis Detection Kit I* (BD Biosciences). In the case of normal mouse colonic tissues [21, 59], cells were incubated with mouse IgGs (5 mg/ml; Innovative Research, IR-MS-GF) at a concentration of 3-5 × 10^5^ cells/100 μl, in order to “block” mouse *Fc receptors* (FcRs) and therefore minimize unspecific binding of antibodies on mouse cells. Cells were subsequently washed and stained with antibodies at dilutions determined by appropriate titration experiments. The antibodies used to visualize distinct sub-types of mouse colon epithelial cells, such as *bottom-of-the-crypt* cells enriched in Lgr5^+^ and Fgfbp1^+^stem/progenitor cells (CBCs; Epcam^+^, Cd44^+^, Kit^neg^), *bottom-of-the-crypt* cells enriched in *goblet cells* (Epcam^+^, Cd44^+^, Kit^+^) and *top-of-the-crypt* cells enriched in *enterocytes* (Epcam^+^, CD44^neg^, CD66a^high^), included: anti-mouse-Epcam-FITC (clone G8.8; Biolegend), anti-mouse-Cd66a-APC/Cy7 (clone Mab-CC1; Biolegend), anti-mouse-Cd44-APC (clone IM7; Biolegend) and anti-mouse-Kit/Cd117-PE (clone 2B8; Biolegend). Mouse stromal cells, intended as cells expressing non-epithelial lineage markers (Stromal-lineage^+^), were excluded by staining with: anti-mouse-Cd3-PE/Cy5 (clone 17A2; BD Biosciences), anti-mouse-Cd16/Cd32-biotin (clone 2.4G2; BD Biosciences), anti-mouse-Cd31-biotin (clone 390; eBioscience) and anti-mouse-Cd45-biotin (clone 30F-11; BD Biosciences). Cells stained with biotin-conjugated antibodies were visualized by secondary staining with Streptavidin-PE-Cy5 (1:200; BD Biosciences). In all experiments, after incubation for 30 min on ice (4°C), stained cells were washed of excess unbound antibodies and resuspended in HBSS with HICS (2%), EDTA (5 mM), penicillin (120 μg/ml), streptomycin (100 μg/ml), amphotericin-B (0.25 μg/ml), HEPES (20 mM), sodium pyruvate (1 mM) and DAPI dilactate (1.1 μM; Molecular Probes). Flow-cytometry analysis was performed using BD FACSAria-II and BD FACSAria-III cell-sorters (BD Biosciences). *Forward-scatter height* versus *forward-scatter width* (FSC-H vs. FSC-W) and *side-scatter height* versus *side-scatter width* (SSC-H vs. SSC-W) profiles were used to eliminate cell doublets. Dead cells were eliminated by excluding DAPI^+^ cells, whereas stromal cells were eliminated by excluding either PacificBlue^+^ cells (in experiments on human CRC xenografts), or PE-Cy5^+^ cells (in experiments on mouse colonic tissues). In cell-sorting experiments, cells were sorted twice sequentially, to achieve higher purity (nozzle diameter: 100 μm). A visual illustration of the gating strategy implemented to sort distinct sub-types of mouse colon epithelial cells is provides as part of **Supplementary Figure 2**. Differences in the mean percentage of Annexin-V^+^ cancer cells between different experimental conditions were tested for statistical significance using two tests: 1) a Welch ANOVA, followed by Dunnett’s T3 test for pairwise comparisons [60]; and 2) a two-way ANOVA test for interaction (cell phenotype vs. chemotherapy). Increases in the percentage of Annexin-V^+^ cancer cells caused by treatment with irinotecan (between paired sets of cancer cells of the same phenotype) were tested for statistical significance using a one-tailed Mann-Whitney U-test.

### Microarray analysis of gene-expression profiles

Microarray profiling of gene-expression patterns was conducted on paired populations of *bottom-of-the-crypt* (EpCAM^+^, CD44^+^, CD166^+^) and *top-of-the-crypt* (EpCAM^+^, CD44^neg^, CD166^neg^) human CRC cells (150,000 cells/replicate), sorted in parallel from 4 independent s.c. tumors representative of the same PDX line (PDX-COLON-8). Of these 4 tumors, 2 originated from animals treated with irinotecan and 2 from animals treated with saline solution. The experimental dataset, therefore, consisted of a total of 8 samples, and included 4 replicates for each of the two main orthogonal variables (cell phenotype vs. *in vivo* treatment). In all 8 samples, the purity of sorted cells after two sequential rounds of FACS was >99%. Sorted cells were flash-frozen in liquid nitrogen, and RNA was purified using the *PureLink™ RNA Micro Scale Kit* (Invitrogen). RNA was amplified and biotinylated using the *GeneChip 3’IVT Express Kit* (Affymetrix) and then hybridized on a *GeneChip®* cartridge containing the *PrimeView™ Human Gene Expression Array* (Affymetrix; platform: GPL15207), following standard protocols recommended by the manufacturer. *GeneChips* were washed and stained in the *Affymetrix Fluidics Station 450*. Gene-expression measurements (i.e., fluorescent signal intensities) were normalized using the *Robust Multi-chip Average* (RMA) method [61] with quantile normalization [62] across multiple arrays. In the case of genes for which multiple oligonucleotide probes were present in parallel within the same array, the aggregate gene-expression levels were summarized by calculating the mean across all relevant probes, after exclusion of “promiscuous” probes (i.e., probes that mapped to more than one gene in the *Entrez* database; https://www.ncbi.nlm.nih.gov/gene). Finally, gene-expression levels were transformed to their log2 values before downstream analyses. The full microarray dataset generated as part of this experiment has been deposited in the *Gene Expression Omnibus* (GEO) repository (https://www.ncbi.nlm.nih.gov/geo) of the *National Center for Biotechnology Information* (NCBI) and is publicly available, under accession number: **GSE213103**.

### Linear modeling of chemotherapy-induced modulation of gene-expression levels

To identify genes that were differentially modulated by *irinotecan* (CPT-11) between colon cancer cells with *bottom-of-the-crypt* (EpCAM^+^, CD44^+^, CD166^+^) and *top-of-the-crypt* (EpCAM^+^, CD44^neg^, CD166^neg^) phenotypes in the PDX-COLON-8 line, we applied a linear model [19, 63–65], using the *linear modeling* (lm) function of the *statistics* (stat) package of the R software (https://www.r-project.org/). The model can be summarized as follows:

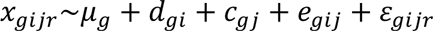

The model envisioned contributions to gene-expression levels that are dependent on the identity of individual genes (*g = individual gene*), the effects of chemotherapy-induced perturbations (*i = no-treatment vs. irinotecan*), the phenotype of analyzed cells (*j = bottom-of-the-crypt* vs. *top-of-the-crypt*) and stochastic effects (r = *experimental replicate*). In this model: *x*_*gijr*_= expression level of gene *g* as measured in the experiment; *μ_g_* = baseline expression level of gene *g*; *d*_*gi*_ = effect of chemotherapy on the expression level of gene *g*; *c*_*gi*_ = effect of cell phenotype on the expression level of gene *g*; *e*_*gij*_ = interaction effect between chemotherapy and cell-phenotype on the expression level of gene *g*; *ε_gijr_* = experimental noise. For each gene, the parameter *e_gij_* captured the degree by which irinotecan induced a preferential change of the gene’s expression in cells with either a *“bottom-of-the-crypt”* (EpCAM^+^, CD44^+^, CD166^+^) or a *“top-of-the-crypt”* (EpCAM^+^, CD44^neg^, CD166^neg^) phenotype. Genes were then ranked based on the value of the *e_gij_* parameter, and those found either above the top 5^th^ percentile (i.e., genes displaying either a higher induction or lower suppression in cells with a *bottom-of-the-crypt* phenotype) or below the bottom 5^th^ percentile (i.e., genes displaying either a higher induction or lower suppression in cells with a *top-of-the-crypt* phenotype) were identified as “responsive” (i.e., differentially modulated by *irinotecan* across different cell types).

### Pathway-enrichment and *Transcription Factor Target Gene* (TFTG) enrichment analysis

To gain insight into the molecular mechanisms responsible for the differential response to irinotecan displayed by different cell-types, the list of genes identified as responsive (i.e., top 5% + bottom 5% of genes, after ranking based on the value of the *e_gij_* interaction parameter) was tested for associations with either (i) genes encoding for mediators of specific biochemical or signaling pathways or (ii) transcriptional targets of known *transcription factors* (TFs). The statistical significance of the association was evaluated using a Fisher exact test (two-tailed), in which the list of genes identified as *“responsive”* was tested for a statistically significant enrichment in genes that belong to specific functional families (i.e., mediators of a specific signaling pathway, transcriptional targets of a specific TF), in the absence of assumptions regarding the magnitude and directionality of the responses, in order to capture signaling pathways and TFs with either preferential induction or lower suppression in either cell type [19]. For each of the pathways and/or TFs identified as associated with a differential response to irinotecan, the magnitude of the association was quantified by computing the *odds ratios* (ORs) for the overlap between gene-lists (i.e., between the list of genes identified as responsive and the list of genes identified as being either mediators of a specific pathway or transcriptional targets of a specific TF). In the specific case of analyses conducted on signaling pathways, the ORs for the enrichment in responsive genes were first calculated considering all responsive genes (i.e., irrespective of the direction of their preferential response, in terms of the cell-type in which they displayed a preferential induction or lower suppression) and then considering, alternatively, either only genes that displayed a higher induction or lower suppression in *bottom-of-the-crypt* (EpCAM^+^, CD44^+^, CD166^+^) cells (i.e., top 5% after ranking based on the interaction parameter), or only genes that displayed a higher induction or lower suppression in *top-of-the-crypt* (EpCAM^+^, CD44^neg^, CD166^neg^) cells (i.e., bottom 5% after ranking based on the interaction parameter), in order to infer the presence of a directionality in the cell-specificity of the effect (i.e., to understand which cell-type displayed preferential induction or lower suppression of a specific signaling pathway). The lists of genes encoding for mediators of specific pathways were derived from the *Molecular Signatures Database* (MSigDB) v4.0 (https://www.gsea-msigdb.org) [66], which incorporates gene signature from both the *Reactome knowledgebase* [67] and the *Pathway Interaction Database* (PID) [68]. The lists of genes representing transcriptional targets of specific TFs have been previously published as part of the *Transcription Factor Target Gene* (TFTG) compendium [18, 69], which was compiled by aggregating published data from *chromatin immuno-precipitation sequencing* (ChIP-seq) experiments and bioinformatics studies on regulatory networks. A list of all the TFs included in the TFTG compendium, and of all their transcriptional targets, is publicly available on the *SimTK* repository (https://simtk.org/projects/tfanno). Signaling pathways identified as associated with a differential response to irinotecan were ranked based on the p-value of the corresponding Fisher exact test, and evaluated for statistical significance after correction for multiple comparisons, using the Benjamini-Hochberg procedure to calculate q-values and estimate *false discovery rates* (FDRs) [70]. The list of signaling pathways identified as differentially responsive to irinotecan (CPT-11) between cells with a *“bottom-of-the-crypt”* (EpCAM^+^, CD44^+^, CD166^+^) and a *“top-of-the-crypt”* (EpCAM^+^, CD44^neg^, CD166^neg^) phenotype is provided in **Supplementary Table 2**. TFs associated with differential response to irinotecan were visualized using a volcano-plot [71], in which each TF included in the TFTG compendium was plotted based on two parameters: 1) the magnitude of the OR describing the overlap between the list of the individual TF’s transcriptional targets and that of genes identified as differentially responsive to irinotecan; and 2) the p-value of the corresponding Fisher exact test (**Figure 1D**). The cutoff chosen for statistical significance of the Fisher exact test used to rank TFs was p<0.00001 (p<10^-5^), in order to provide a conservative Bonferroni correction for multiple comparisons (total number of TFs tested in parallel: < 2,000). In order to visualize the directionality of the transcriptional responses identified as dependent on TFs with differential activation between populations (i.e., E2F4, TFDP1), all genes for which a measurement in expression was available were plotted on a two-axis graph, based on their response to irinotecan (i.e., average fold-change in mean expression levels between treated and untreated samples) in *“top-of-the-crypt”* (EpCAM^+^, CD44^neg^, CD166^neg^; x-axis) and *“bottom-of-the-crypt”* (EpCAM^+^, CD44^+^, CD166^+^; y-axis) cancer cells, after stratification of the genes into *“targets”* and *“non-targets”* of the corresponding TF **(Supplementary Table 3, Figure 1E**). In both cases (E2F4, TFDP1) the visual exploration revealed a preferential up-regulation of target genes in *“top-of-the-crypt”* (EpCAM^+^, CD44^neg^, CD166^neg^) cancer cells, which are *non-tumorigenic* (NT), as compared to *“bottom-of-the-crypt”* (EpCAM^+^, CD44^+^, CD166^+^) cancer cells, which are enriched in cells with a *“cancer stem cell”* (CSC) phenotype. In order to quantify the magnitude of the difference between the two cell-types in such response to irinotecan (i.e., up-regulation of E2F4 and TFDP1 *target* genes), we computed average fold-changes in gene expression levels for *target* and *non-target* genes across the two cell-types, as well as the average difference in such fold-changes between the two cell-types, and then plotted such averages side-by-side, alongside their 95% *confidence intervals* (**Figure 1F**). Finally, differences between such average fold-changes in gene expression were tested for statistical significance (Welch’s t-test, two-tailed, assuming unequal variance).

### Bioinformatic analysis of *E2F4* and *TFDP1* mRNA expression levels in the *colon adenocarcinoma* (COAD) and *rectal adenocarcinoma* (READ) datasets of *The Cancer Genome Atlas* (TCGA) public repository

To understand whether the expression levels of *E2F4* and *TFDP1* displayed systematic differences between normal human colorectal tissues and human primary CRCs, we studied the *colon adenocarcinoma* (COAD) and *rectal adenocarcinoma* (READ) datasets of *The Cancer Genome Atlas* (TCGA) public repository (https://portal.gdc.cancer.gov; downloaded: February 16, 2017) [72, 73]. *RNA-sequencing* (RNA-seq) experiments from the two datasets (COAD, READ) were combined into a single dataset, after exclusion of all experiments considered redundant and/or duplicative (i.e., experiments performed on multiple tumor specimens collected from the same patient, in which case we retained only the experiment performed on the first specimen collected from the primary tumor). The final dataset consisted of RNA-seq data representative of 51 normal colon and rectal tissues (COAD: n=41; READ: n=10; male: n=23; female: n=28) and 622 CRCs (COAD: n=456; READ: n=166; male: n=330; female: n=289; unknown sex: n=3). The metric adopted to quantify mRNA expression levels was the log_2_ of the number of FPKMs (*fragments per kilobase of exon per million mapped reads*). The presence of a correlation between *E2F4* and *TFDP1* expression levels was visualized using a scatter-plot and quantified by computing a Person’s correlation coefficient (r) between the FPKM values for the corresponding mRNA species, across the full dataset. The statistical significance of the Pearson’s correlation coefficient (r) was evaluated using a two-tailed t-test for correlation coefficients (null hypothesis: r=0). For each of the two analyzed genes (*E2F4*, *TFDP1*), the distribution of mRNA expression levels across the two sample types (normal vs. cancer) was visualized using violin-plots [74] with staggered data-points. Violin plots were superimposed to box-plots [75] to allow for better visualization of each distribution’s median and quartiles. Differences in the distribution of log_2_ FPKM values between normal colonic tissues (n=51) and CRCs (n=622) were tested for statistical significance using a Mann-Whitney U-test (two-tailed).

### RNA-seq analysis of *E2f4* and *Tfdp1* mRNA expression across different cellular lineages of the mouse normal colonic epithelium

The expression levels of *E2f4* and *Tfdp1* mRNAs were evaluated for differences between 3 different sub-types of mouse colon epithelial cells, including: 1) *bottom-of-the-crypt* cells enriched in Lgr5^+^ and Fgfbp1^+^ stem/progenitor cells (Epcam^+^/Cd44^+^/Cd66a^low^/Kit^neg^); 2) *bottom-of-the-crypt* cells enriched in *goblet cells* (Epcam^+^/Cd44^+^/Cd66a^low^/Kit^+^); 3) and *top-of-the-crypt* cells enriched in terminally differentiated *enterocytes* (Epcam^+^/Cd44^neg^/CD66a^high^). The analysis was conducted by performing RNA-seq on purified preparations of the three cell types, isolated in parallel by FACS from the same set of primary colonic tissues (**Figure 2A, Supplementary Figure 2**), following established protocols [21, 59]. Briefly, for each of the 3 cell-types, 3 biological replicates were generated, starting from 3 independent adult mice (6-8 weeks old, female) of the C57BL/6J strain (The Jackson Laboratory; stock #000664). Sorted cells were flash-frozen in liquid nitrogen immediately after purification by FACS. Total RNA was extracted from each sample using the *NuceloSpin RNA XS* kit (Macherey-Nagel) and analyzed for yield and integrity using a *Bioanalyzer* instrument (Agilent), which enabled calculation of *RNA integrity numbers* (RINs) [76]. All RNA specimens used for RNA-seq experiments were confirmed to be of high quality (RIN>8) [77]. RNA-seq experiments were performed with the assistance of the *J.P. Sulzberger Columbia Genome Center*. Briefly, RNA-seq libraries were generated using the *Illumina Stranded mRNA Prep* kit, which employs poly-A oligonucleotides to selectively “pull-down” *messenger RNAs* (mRNAs). RNA-seq libraries were sequenced using either an *Illumina HiSeq 4000* machine (30 million, single-end, 100 bp reads) or an *Illumina NovaSeq 6000* machine (20 million, paired-end, 100 bp reads). RNA-seq data were mapped onto the *University of California Santa Cruz* (UCSC) mm10 mouse genome using the *OLego* software [78], and then normalized using the *Cufflinks* computational pipeline [79, 80]. Gene-expression levels were quantified by computing the log2 of *Transcripts per Million* (TPM) values. Differences in mean log_2_ TPM values across different populations were tested for statistical significance using a one-way ANOVA test for repeated measures. The raw sequencing data (FASTQ) generated by this RNA-seq experiment have been deposited in the NCBI-GEO repository (https://www.ncbi.nlm.nih.gov/geo) and are publicly available, under accession number: **GSE202483**.

### Cell lines

The two CRC cell lines used in this study are available from the *American Type Culture Collection* (ATCC; http://www.atcc.org) and include: HCT116 cells (ATCC catalog: CCL-247) [81] and HT29 cells (ATCC catalog: #HTB-38) [82, 83]. Lentivirus vectors were *“packaged”* using HEK293 human embryonic kidney cells (GenHunter; catalog: Q401). All cell lines were cultured in RPMI-1640 tissue-culture medium (Sigma-Aldrich), supplemented with 10% *heat-inactivated fetal bovine serum* (HI-FBS), L-glutamine (2 mM), penicillin (120 μg/ml), streptomycin (100 μg/ml), HEPES (20 mM; Sigma-Aldrich) and sodium pyruvate (1 mM; Lonza).

### Lentivirus vectors encoding *short-hairpin RNA* (shRNA) constructs for the suppression of E2F4 expression

The sequence and targeting sites of the two shRNA constructs used in this study to knock-down E2F4 expression (TRCN0000013808, TRCN0000013809) are described in **Supplementary Table 4**. Both shRNA constructs were developed by *The RNAi Consortium* (TRC) of the *Broad Institute* [25] and are currently available from *Horizon Discovery* (Perkin Elmer). The two shRNA constructs were cloned into the *pSIH1-H1-copGFP* lentivirus backbone (*System Biosciences;* catalog: #SI501A-1), which carries the *copepod green fluorescent protein* (copGFP) as a green, fluorescent reporter (isolated from the marine crustacean *Pontellina plumata*). Lentivirus infectious particles were produced following established protocols and procedures [84], using a 3^rd^ generation lentivirus vector system [85, 86] with minor methodological improvements [9, 58]. Very briefly, lentivirus infectious particles were produced by co-transfection in HEK293 human embryonic kidney cells (GenHunter; catalog: Q401) of four independent plasmids, including three plasmids encoding for distinct structural and/or functional elements of the virion (pMDLg/pRRE, Addgene #12251; pRSV-Rev, Addgene #12253; pCMV-VSV-G, Addgene #8454; https://www.addgene.org) [86], and one plasmid encoding for the provirus (*pSIH1-H1-copGFP* +/-shRNAs targeting *E2F4*), designed to include three key features: 1) a modified 5’ *long terminal repeat* (LTR) sequence, in which the U3 region was replaced by the *Respiratory Syncytial Virus* (RSV) promoter in order to make the lentivirus packaging Tat-independent [87, 88]; 2) a modified 3’ *long terminal repeat* (LTR) sequence in which the U3 region was deleted, in order to render it *self-inactivating* (SIN) upon integration in the host’s genomic DNA [87, 88]; and 3) the sequence of the shRNA transgene of interest (i.e., one of the shRNAs targeting *E2F4*) under the transcriptional control of the H1 promoter (RNA Polymerase-III) [89]. Plasmids were transfected into HEK293 cells by precipitation in *calcium-phosphate* (Ca-P), following standard protocols [90]. HEK293 cells were then incubated in tissue-culture media supplemented with caffeine (4 mM) to increase the yield of lentivirus infectious particles in cell supernatants [91], which were harvested 24-48 hours after the end of the transfection procedure and immediately filtered to remove cellular debris (filter pore size: 0.45 μm). Lentivirus infectious particles were concentrated by ultra-centrifugation (70,000 g, 2 hours at 4°C), in order to achieve high titers and enable high *multiplicity of infection* (MOI). Concentrated viral particles (MOI>10) were used to infect single-cell suspensions of CRC cells by *“spinoculation”* (i.e., centrifugation at 1,200 g, 2 hours, 4 °C) in the presence of polybrene (8 μg/ml), followed by incubation at 37 °C for 2 hours [92, 93]. Infected cells were subsequently propagated either as *two-dimensional* (2D) monolayers (HT29, HCT116) or *three-dimensional* (3D) organoids (PDX-COLON-8, PDX-COLON-60, PDX-COLON-136, PDX-COLON-441) and analyzed by flow cytometry 72 hours after infection, to assess infection efficiency. In all experiments designed to either establish the capacity of shRNA constructs to either suppress E2F4 expression or evaluate its functional effects, analyses were conducted on purified preparations of human genetically engineered CRC cells, sorted by FACS based on simultaneous expression of the lentivirus-encoded fluorescent reporter (copGFP^+^) and human epithelial markers characteristic of a *bottom-of-the-crypt* phenotype (EpCAM^+^, CD44^+^, CD166^+^).

### *Three-dimensional* (3D) *in vitro* organoid cultures

In selected experiments, human CRC cells isolated from PDX lines were propagated *in vitro* as 3D organoids, following previously described tissue-culture protocols [94, 95], with minor modifications. Briefly, human CRC cells were purified by FACS, infected with lentivirus vectors, resuspended in a 1:1 mixture of *Dulbecco’s Modified Eagles’ Medium* and *Ham’s F12* medium (DMEM/F12), supplemented with mouse Wnt3a (100 ng/ml; Peprotech), human RSPO1 (1 μg/ml; R&D), human EGF (50 ng/ml; Invitrogen), human NOGGIN (100 ng/ml; Peprotech), Y-27632 (10 μM; Calbiochem), nicotinamide (10 mM; Sigma), N-Acetyl-L-cysteine (1 mM; Sigma), *B-27 Supplement Without Vitamin-A* (Invitrogen) and *N-2 Supplement* (Invitrogen), and finally plated on top of a layer of *Growth Factor Reduced* (GFR) *Basement Membrane Matrigel* (Corning), supplemented with the Jagged-1 (188-204) peptide (1 uM; Peprotech). In experiments designed to compare the effects of different shRNA constructs on the organoid-forming capacity of cancer cells, human lentivirus-infected malignant cells with a *bottom-of-the-crypt* phenotype (copGFP^+^, EpCAM^+^, CD44^+^, CD166^+^) were purified by FACS, and seeded at equal numbers in parallel replicates within multi-well plates (2,000 cells/well). The number of 3D organoids formed upon plating of sorted cells was counted 3 weeks after seeding (with organoids defined as viable 3D multicellular structure consisting of >50 cells, as estimated visually). Differences in the number of organoids formed by cells engineered with different shRNAs were tested for statistical significance using a Welch ANOVA test, followed by Dunnett’s T3 test for pairwise comparisons [60].

### Analysis of *E2F4* mRNA expression levels by *reverse-transcription and quantitative polymerase chain-reaction* (RT-qPCR)

The down-regulation of *E2F4* mRNA expression following infection with lentivirus vectors encoding for shRNA constructs targeting *E2F4* was evaluated by RT-qPCR, based on the analytical procedure suggested by Kenneth J. Livak and Thomas D. Schmittgen [96]. Briefly, total RNA was extracted from lentivirus-infected cells using TRIzol (Invitrogen) [97, 98] and then precipitated using glycogen (Invitrogen). Reverse-transcription and cDNA synthesis were performed using the *SuperScript-III* reverse transcription kit (Invitrogen). RT-qPCR reactions were run on an ABI-7900HT *Fast Real-Time PCR System* (Applied Biosystems), using a TaqMan™ assay specific for the *E2F4* mRNA (probe: Hs00608098_m1; Applied Biosystems), following previously published protocols [59]. Differences in *E2F4* mRNA expression levels between reference samples (e.g., cells infected with empty lentivirus vectors) and tested specimens (e.g. cells infected with lentivirus vectors encoding for shRNA constructs targeting *E2F4*) were measured as fold-changes, calculated based on delta-delta Ct (ΔΔCt) values (ΔCt[empty vector]-ΔCt[shRNA]), after normalization to *ACTB* (*β-Actin*) mRNA expression levels (probe: Hs00357333_g1; Applied Biosystems) [96]. Finally, differences in *E2F4* mRNA expression levels between cells infected with different types of lentivirus constructs were tested for statistical significance using a two-way ANOVA test for interactions (cell line vs. lentivirus construct) on ΔΔCt values (null hypothesis: ΔCt[empty vector]-ΔCt[shRNA]=0; fold-change=1) followed by a Dunnett’s test on pre-selected pair-wise comparisons.

### Western blot

The down-regulation of E2F4 protein expression levels following infection with lentivirus vectors encoding for shRNA constructs was evaluated by Western blotting, performed according to previously described protocols and procedures [59, 99, 100]. Briefly, lentivirus-infected cells were first soaked in a hypotonic buffer, containing Tris-HCl at pH=7.5 (10mM), MgCl_2_ (5 mM) and *cOmplete™ Proteinase Inhibitor Cocktail* (Roche), for 20 min on ice (4°C). Cells were then lysed in *sodium dodecyl-sulfate* (SDS) *Sample Loading Buffer*, containing Tris-HCl with pH=6.8 (50 mM), SDS (2% w/v), glycerol (10% w/v), EDTA (5 mM), bromophenol blue (0.02% w/v) and β-mercapto-ethanol (3% w/v). Protein extracts were separated by denaturing *SDS-polyacrylamide gel electrophoresis* (SDS-PAGE) and electro-blotted onto a *poly-vinylidene-difluoride* (PVDF) membrane (Amersham). PDVF membranes were incubated with a *“blocking”* solution, consisting of DPBS containing skim-milk (5% w/v) and Tween-20 (0.05%), in order to saturate unspecific protein binding sites (i.e., prevent unspecific binding by antibodies), and then incubated with primary antibodies, consisting of either a rabbit-anti-human-E2F4 (clone: EPR8259; Abcam) or a mouse-anti-human-β-Actin (clone: AC-15; Millipore Sigma) monoclonal antibody. Primary antibodies were revealed by secondary staining with either goat-anti-rabbit-IgG or sheep-anti-mouse-IgG polyclonal antibodies conjugated to horseradish peroxidase (Amersham). Staining by secondary antibodies was visualized by *enhanced chemiluminescence* (ECL), using the *SuperSignal West Dura Extended Duration Substrate* (Thermo Scientific).

### Evaluation of tumorigenic cell frequency by *Extreme Limiting Dilution Analysis* (ELDA)

To understand whether E2F4 down-regulation would result in a reduction of the frequency of cells with tumor-initiating capacity, we used ELDA [101] to calculate the frequency of tumorigenic cells among cells with a *bottom-of-the-crypt* phenotype (EpCAM^+^, CD44^+^, CD166^+^), which are known to be selectively enriched in cells with *“cancer stem cell”* (CSC) properties [8, 9], following infection with either an *“empty”* lentivirus vector or a lentivirus vector encoding for an shRNA construct targeting E2F4 Very briefly, human lentivirus-infected CRC cells with a *bottom-of-the-crypt* phenotype (copGFP^+^, EpCAM^+^, CD44^+^, CD166^+^) were sorted by FACS, diluted to appropriate injection doses, mixed with BD Matrigel (BD Biosciences) at a 1:1 ratio, and injected sub-cutaneously (s.c). in immunodeficient NSG mice, following previously published protocols [8, 9]. To limit possible confounding effects related to biological differences between individual animal hosts, the two experimental groups undergoing comparisons (i.e., cells infected with empty lentivirus vectors vs. cells infected with lentivirus vectors encoding for shRNA constructs targeting E2F4) were injected as paired sets, whereby equal doses of the two cell preparations were injected into opposite flanks of the same animals. Injected mice were monitored weekly for tumor engraftment up to a maximum of 10 months, and euthanized once s.c. tumors reached a maximum diameter of 20 mm. Tumorigenic cell frequencies were calculated using the computational pipeline developed by Yifang Hu and Gordon K. Smyth to analyze ELDA results, which is publicly available on the website of the *Bioinformatics Division* of the *Walter and Eliza Hall Institute of Medical Research* (http://bioinf.wehi.edu.au/software/elda) [101]. Differences in tumorigenic cell frequency were tested for statistical significance on the same platform, performing a *Likelihood-Ratio Test* (LRT), in which the distribution of the test’s statistic (i.e., a natural logarithm of the likelihood ratio) is approximated using the χ^2^ distribution, based on Wilk’s theorem [101].

### Quantification of apoptosis by immunofluorescence (Caspase-3)

Analysis of copGFP and cleaved Caspase-3 protein expression in human CRC tumor xenografts was performed by *immunofluorescence* (IF) on frozen tissue-sections, following previously reported procedures [102], with minor modifications. Very briefly, tumor tissues were fixed by immersion in a 10% formalin (v/v) phosphate-buffered solution (2 hours at room temperature), then infiltrated with a 30% sucrose solution to limit cryogenic artifacts (2 days, 4°C) and finally embedded in *Optimal Cutting Temperature* (OCT) compound (Sakura Finetek) and stored in a deep-freezer (−80°C), before being cut on a refrigerated cryo-microtome. Shortly before staining, tissue sections were washed twice with DPBS (5 minutes at room temperature) and then incubated (30 minutes at room temperature) with a *“blocking”* solution consisting of DPBS supplemented with Triton X-100 (0.1% w/v), bovine serum albumin (2% w/v) and normal donkey serum (5% w/v), in order to saturate unspecific protein binding sites (i.e., limit the unspecific binding of antibodies). Tissue sections were then stained (12 hours, 4°C) with a rabbit-anti-human-cleaved-Caspase-3 (cleavage site: Asp175) polyclonal antibody (Cell Signaling; catalog: #9661). Stained tissue-sections were washed three times with DPBS containing Triton X-100 (0.1% w/v) to remove excess unbound primary antibodies, and then stained (1 hour, 4°C) with an affinity-purified donkey-anti-rabbit-IgG (H+L) polyclonal antibody conjugated to Alexa Fluor 594 (Jackson ImmunoResearch). Finally, stained tissue-sections were washed three times with DPBS containing Triton X-100 (0.1% w/v) to remove excess unbound secondary antibodies, then mounted using the *ProLong^®^ Gold Antifade Reagent with DAPI* (Molecular Probes) and imaged using a Zeiss LSM-700 scanning confocal microscope (Carl Zeiss). The percentage of apoptotic tumor cells was computed manually, dividing the number of malignant cells displaying expression of cleaved Caspase-3 (numerator: n. of cells with copGFP^+^/Caspase-3^+^ phenotype) by the total number of malignant cells (denominator: n. of cells with copGFP^+^ phenotype) across 10-21 independent fields of the same size, each representing an experimental replicate. Differences in the percentage of cancer cells displaying expression of cleaved Caspase-3 between different experimental conditions were tested for statistical significance using three different tests: i) a Kruskal-Wallis H-test, followed by a Dunn’s test for pre-selected pair-wise multiple comparisons; ii) a Welch ANOVA test (assuming unequal variance), followed by a Dunnett’s T3 test for pre-selected pair-wise comparisons [60]; and iii) a 2-way ANOVA test for interaction between type of lentivirus vector and chemotherapy, followed by a Dunnett’s test for pre-selected pair-wise comparisons.

## COMPETING FINANCIAL INTERESTS

|  |  |
| --- | --- |
| **Junichi Matsubara:** | No competing interests to disclose |
| **Yong Fuga Li:** | Yong Fuga Li is currently an employee of **Illumina Inc.** |
| **Sanjay Koul:** | No competing interests to disclose |
| **Junko Mukohyama:** | No competing interests to disclose |
| **Luis Enrique Valencia Salazar:** | No competing interests to disclose |
| **Taichi Isobe:** | No competing interests to disclose |
| **Dalong Qian:** | No competing interests to disclose |
| **Michael F. Clarke:** | Michael F. Clarke is a founder of **Tachyon Therapeutics**. Michael F. Clarke is listed as a co-inventor on patents and patent applications owned by the *University of Michigan* (**US-07723112-B2**) and *Stanford University* (**US-09329170-B2**, **US-09850483-B2**), which resulted in the award of royalties and/or stock from **Oncomed Pharmaceuticals** and **Quanticel Pharmaceuticals**. |
| **Debashis Sahoo:** | No competing interests to disclose |
| **Russ B. Altman:** | No competing interests to disclose |
| **Piero Dalerba:** | Piero Dalerba is listed as a co-inventor on patents and patent applications owned by the *University of Michigan* (**US-07723112-B2**), *Stanford University* (**US-09329170-B2**, **US-09850483-B2**, **US-10344094-B2**, **US-11130813-B2**, **US-12037402-B2**) and *Columbia University* (**US-12115140-B2**), which resulted in the award of royalties and/or stock from **Oncomed Pharmaceuticals**, **Quanticel Pharmaceuticals** and **Forty Seven Inc. (Gilead)**. Piero Dalerba's spouse is employed by **Regeneron Pharmaceuticals** and owns stock of the following pharmaceutical companies: **AbbVie**, **Amgen**, **AstraZeneca**, **Eli Lilly and Company**, **GlaxoSmithKline**, **Johnson & Johnson**, **Merck & Co**, **Novartis**, **Organon**, **Pfizer** and **Viatis**. |

## AUTHOR CONTRIBUTIONS (CRediT)

|  |  |
| --- | --- |
| **Junichi Matsubara:** | Conceptualization, Methodology, Investigation, Visualization, Formal analysis, Data curation, Funding acquisition, Validation, Writing (original draft), Writing (review and editing) |
| **Yong Fuga Li:** | Conceptualization, Methodology, Investigation, Visualization, Formal analysis, Software, Data curation, Validation, Writing (original draft), Writing (review and editing) |
| **Sanjay Koul:** | Conceptualization, Methodology, Investigation, Visualization, Formal analysis, Data curation, Validation, Writing (original draft), Writing (review and editing) |
| **Junko Mukohyama:** | Conceptualization, Methodology, Investigation, Visualization, Formal analysis, Data curation, Validation, Writing (original draft) |
| **Luis Enrique Valencia Salazar:** | Methodology, Investigation, Visualization, Formal analysis, Data curation, Validation |
| **Taichi Isobe:** | Investigation, Formal analysis, Data curation, Validation |
| **Dalong Qian:** | Investigation, Validation |
| **Michael F. Clarke:** | Conceptualization, Methodology, Funding acquisition, Project administration, Validation, Supervision, Resources, Writing (original draft) |
| **Debashis Sahoo:** | Conceptualization, Methodology, Visualization, Formal analysis, Software, Data curation, Funding acquisition, Writing (review and editing) |
| **Russ B. Altman:** | Conceptualization, Methodology, Visualization, Formal analysis, Data curation, Funding acquisition, Project Administration, Supervision, Resources, Writing (original draft), Writing (review and editing) |
| **Piero Dalerba:** | Conceptualization, Methodology, Investigation, Visualization, Formal analysis, Data curation, Funding acquisition, Project administration, Supervision, Resources, Validation, Writing (original draft), Writing (review and editing) |

## ABBREVIATIONS

ADC: antibody-drug conjugate
CD166/ALCAM: cluster of differentiation 166 antigen/activated leukocyte cell adhesion molecule
CD44: cluster of differentiation 44 antigen
CPT: camptothecin
CPT-11: 7-ethyl-10-[4-(1-piperidino)-1-piperidino]carbonyloxy-camptothecin (irinotecan)
copGFP: copepod green fluorescence protein (Pontellina plumata)
CRC: colorectal carcinoma
CSCs: cancer stem cells
DREAM: dimerization partner (DP), retinoblastoma-like (RB-like), E2F and multi-vulva class B (MuvB) complex
DSB: double-strand DNA break
DYRK1A: dual specificity tyrosine phosphorylation regulated kinase 1A
E2F4: E2F transcription factor 4;
ELDA: extreme limiting dilution analysis
FACS: fluorescence activated cell sorting
FDR: false discovery rate
EPCAM: epithelial cell adhesion molecule
LIN9: human homolog of the lin-9 gene (Caenorhabditis elegans)
LIN37: human homolog of the lin-37 gene (Caenorhabditis elegans)
LIN52: human homolog of the lin-52 gene (Caenorhabditis elegans)
LIN54: human homolog of the lin-54 gene (Caenorhabditis elegans)
MuvB: multi-vulva class B (MuvB) complex
OR: odds ratio
PDX: patient-derived xenograft
RBBP4: retinoblastoma protein (RB) binding protein 4, chromatin remodeling factor
RBL1: retinoblastoma protein (RB) transcriptional corepressor like 1
RBL2: retinoblastoma protein (RB) transcriptional corepressor like 2
RNA-seq: RNA-sequencing
shRNA: short hairpin RNA
siRNA: short interfering RNA
SN-38: 7-ethyl-10-hydroxycamptothecin
SSB: single-strand DNA break
TFDP1: transcription factor dimerization partner 1
TFTG: transcription factor target genes
TOP1: topoisomerase 1

## ACKNOWLEDGEMENTS

We are grateful to Gary L. Mantalas, Sopheak Sim and Norma F. Neff (*Stanford University*), as well as Erin Bush (*Columbia University*), for their technical guidance in the design and execution of transcriptional profiling experiments, using either gene-expression microarrays or RNA-sequencing technologies. We also wish to thank Jennifer Cheung and Patty Lovelace (*Stanford University*), as well as Michael Kissner (*Columbia University*) for their assistance with *fluorescence activated cell sorting* (FACS) experiments.

---

**Supplementary Figure S1.**
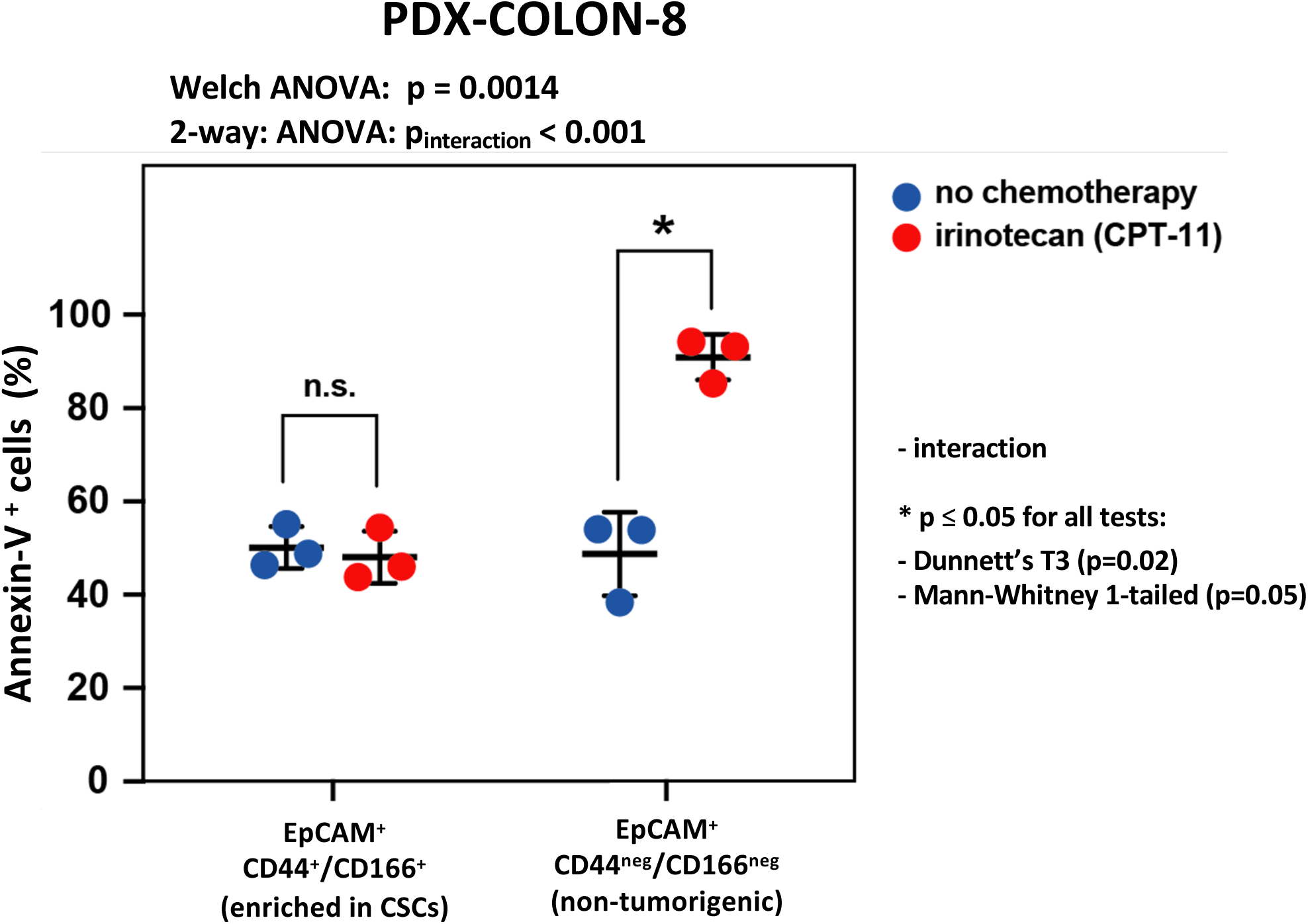
Irinotecan (CPT-11) displays preferential cytotoxicity against the subset of colon cancer cells with a *top-of-the-crypt* phenotype (EpCAM^+^, CD44^neg^, CD166^neg^) as compared to the subset with a *bottom-of-the-crypt* phenotype (EpCAM^+^, CD44^+^, CD166^+^) To understand whether colon cancer cells with a *cancer stem cell* (CSC) phenotype displayed preferential resistance to chemotherapy, we tested whether, in immune-deficient mice engrafted with a human colon cancer *patient-derived xenograft* (PDX) line, *in vivo* treatment with irinotecan (CPT-11) was more capable of inducing apoptosis in cancer cells with a *top-of-the-crypt* (EpCAM^+^, CD44^neg^, CD166^neg^) phenotype, which are non-tumorigenic, as compared to cancer cells with a *bottom-of-the-crypt* phenotype (EpCAM^+^, CD44^+^, CD166^+^), which are enriched in *cancer stem cells* (CSCs). Adult, female, NOD/SCID/IL2Rg^-/-^ (NSG) immune-deficient mice were engrafted sub-cutaneously (s.c.) with the PDX-COLON-8 line, which is known to contain populations with both *top-of-the-crypt* (EpCAM^+^, CD44^neg^, CD166^neg^) and *bottom-of-the-crypt* (EpCAM^+^, CD44^+^, CD166^+^) phenotypes. Tumor-bearing mice were then treated with either irinotecan (50 μg/g, once weekly x 4 weeks, i.p.) or a placebo control (1 ml of saline solution, once weekly x 4 weeks, i.p.) and sub-cutaneous tumors harvested 48 hours after the last treatment (day 24). The percentage of apoptotic cells was quantified by flow cytometry, by measuring the percentage of Annexin-V^+^ cancer cells in each of the two phenotypic sub-populations. *In vivo* treatment with irinotecan (CPT-11) induced preferential apoptosis among cancer cells with a *top-of-the-crypt* phenotype (EpCAM^+^, CD44^neg^, CD166^neg^) as compared to cancer cells with a *bottom-of-the-crypt* phenotype (EpCAM^+^, CD44^+^, CD166^+^). Differences in the percentage of Annexin-V^+^ cancer cells between populations were tested for statistical significance using; 1) a Welch ANOVA across the full dataset (assuming unequal variance), followed by a Dunnett’s T3 test for pre-specified pairwise comparisons; and 2) a two-way ANOVA test for interaction (cell phenotype vs. chemotherapy). Increases in the percentage of Annexin-V^+^ cancer cells caused by treatment with irinotecan were tested for statistical significance using a one-tailed Mann-Whitney U-test. Error bars: mean +/-standard deviation.

**Supplementary Figure S2.**
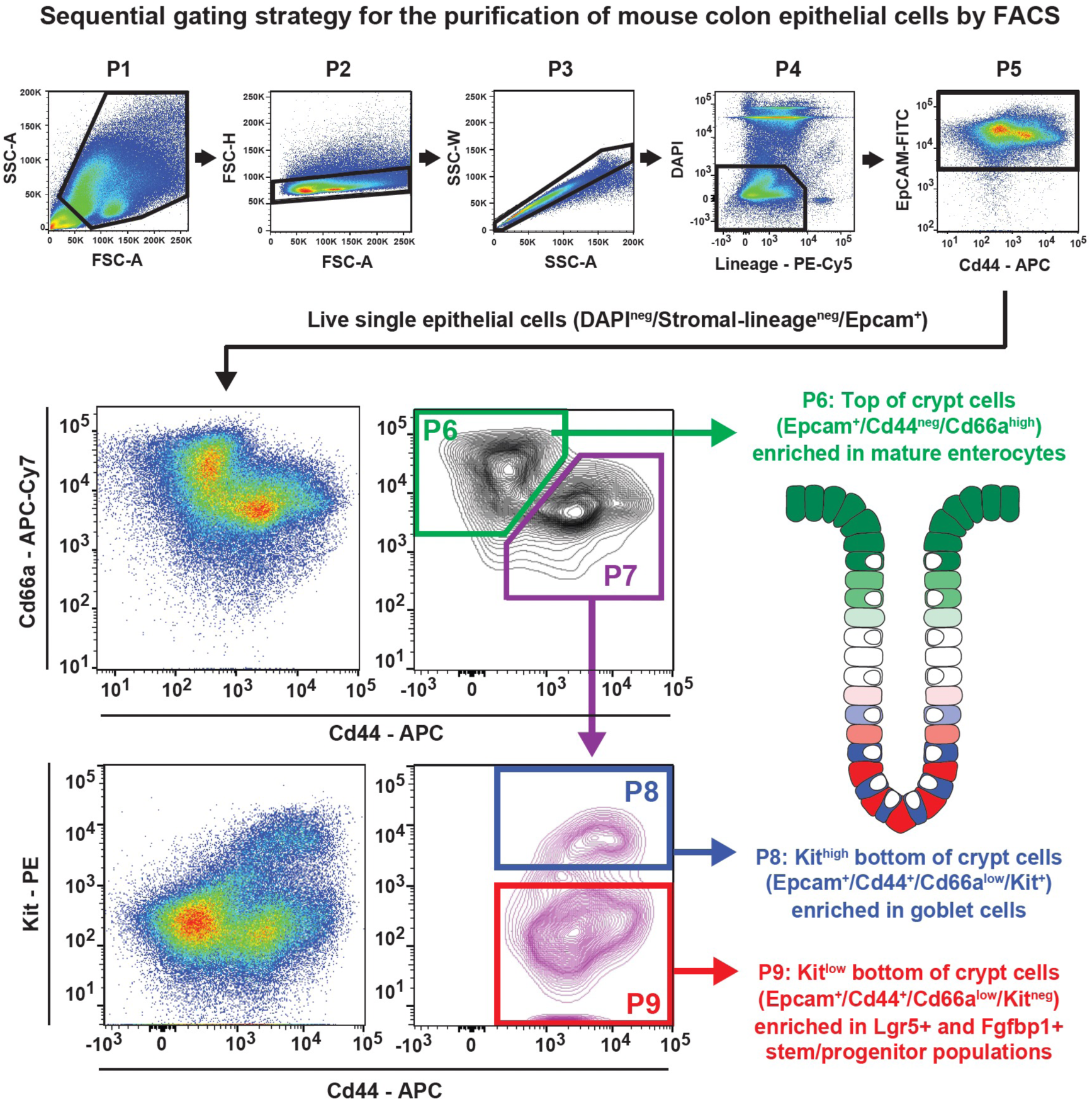
Schematic representation of the gating strategy used for the differential purification of distinct sub-types of colon epithelial cells by *fluorescence-activated cell sorting* (FACS) Single-cell suspensions obtained from the dissociation of mouse colonic crypts were sorted by FACS, using nine sequential gates organized hierarchically, and structured as follows: **(P1)** separation of whole cells from small-sized cellular fragments, based on *forward-scatter area* (FSC-A) and *side-scatter area* (SSC-A) profiles; **(P2)** first separation of single-cells from cell-doublets, based on *forward-scatter area* (FSC-A) vs. *forward-scatter height* (FSC-H) profiles; **(P3)** second separation of single-cells from cell-doublets, based on and *side-scatter area* (SSC-A) vs. *side-scatter width* (SSC-W) profiles; **(P4)** separation of live cells of non-stromal lineage (DAPI^neg^/Lineage^neg^) from dead cells (DAPI^+^) and cells of stromal lineage (Lineage^+^); **(P5)** separation of epithelial cells (EpCAM^+^) from non-epithelial cells (EpCAM^neg^); **(P6-P7)** separation of epithelial cells with a *“top-of-the-crypt”* phenotype, which are enriched in enterocytes (P6: Epcam^+^, CD44^neg^, CD66a^high^), from epithelial cells with a *“bottom-of-the-crypt”* phenotype (P7: Epcam^+^, CD44^+^, CD66a^low^); **(P8-P9)** splitting of epithelial cells with a *“bottom-of-the-crypt”* phenotype into a Kit^+^ sub-group (P8), enriched in goblet cells, and a Kit^neg^ sub-group (P9), enriched in both Lgr5^+^ and Fgfbp1^+^ stem/progenitor cells. Cells belonging to stromal lineages (Lineage^+^) were excluded by staining with a cocktail of antibodies directed against surface markers characteristic of hematopoietic and endothelial lineages (Cd3, Cd45, Cd16, Cd31, Cd32).

**Supplementary Figure S3.**
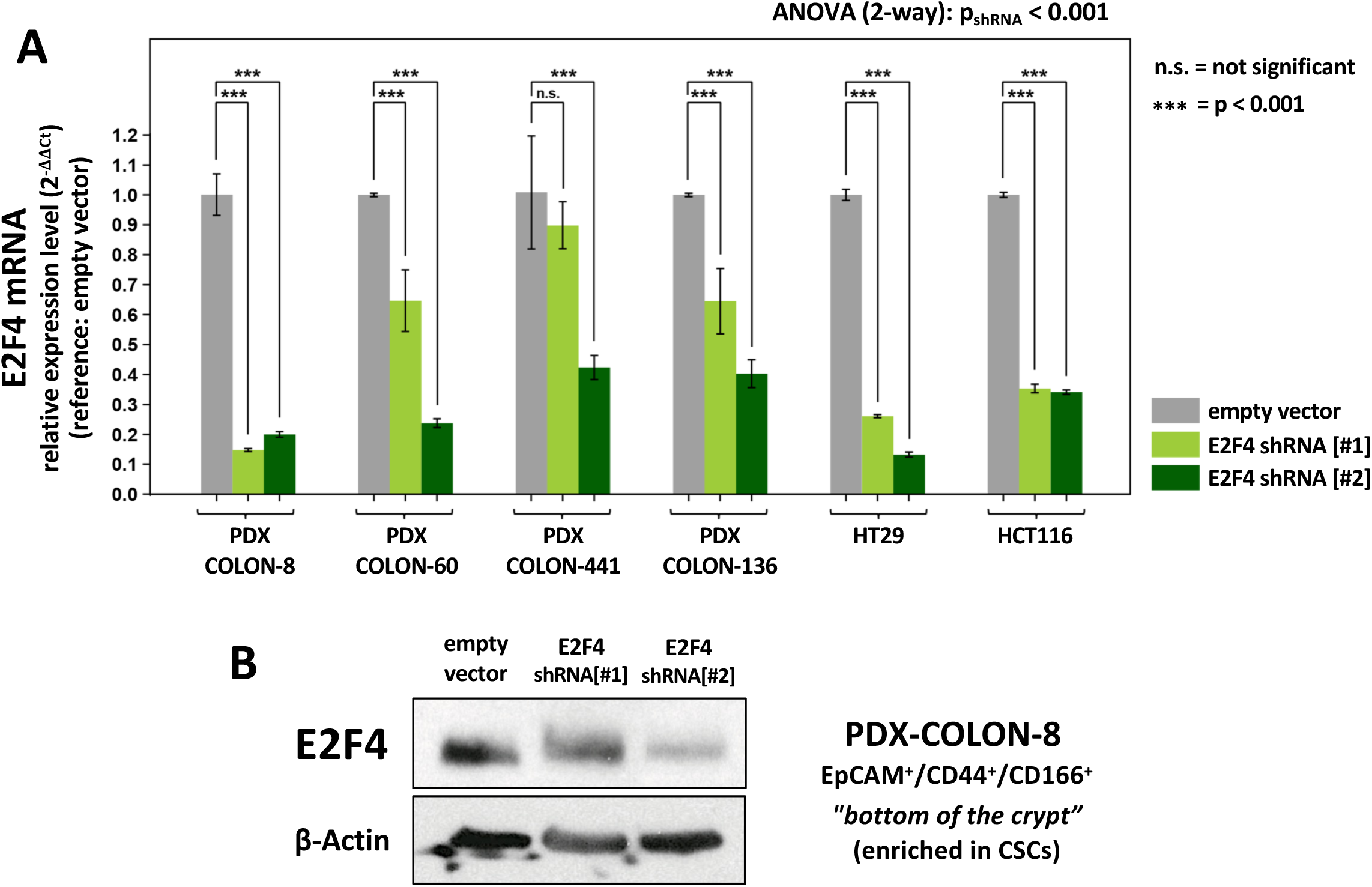
Evaluation of E2F4 knock-down efficiency using *short-hairpin RNA* (shRNA) constructs. **(A)** Analysis by RT-qPCR of *E2F4* mRNA expression in *colorectal cancer* (CRC) cells infected with lentivirus vectors encoding for shRNA constructs targeting the *E2F4* mRNA. The analysis confirmed the capacity of both shRNA constructs used in this study, E2F4-shRNA[#1] and E2F4-shRNA[#2], to cause a reduction in *E2F4* expression levels, as compared to what observed in parental cells infected with an empty vector (reference standard). The analysis was conducted on 6 independent models, including: a) four *patient derived xenograft* (PDX) lines (PDX-COLON-8, PDX-COLON-60, PDX-COLON-C441, PDX-COLON-136); and b) two conventional cell lines grown as *two-dimensional* (2D) monolayers (HT29, HCT116). In both cases, the analysis was conducted on purified preparations of lentivirus-infected malignant cells, isolated by FACS from primary tissues based on the expression of a green fluorescent reporter (copGFP) encoded by the lentivirus construct. Results are reported as fold-changes relative to cells infected with an empty vector, after ΔΔCt normalization to *ACTB* mRNA expression levels and calculation of 2^-ΔΔCt^ values (Livak & Schmittgen, *Methods*, 25:402-408, 2001). Differences in expression levels were tested for statistical significance using a two-way ANOVA test on ΔΔCt values (cell line vs. lentivirus construct) followed by Dunnett’s test on pairwise comparisons. Error bars: mean +/-standard deviation. **(B)** Analysis by Western blot of E2F4 protein expression in cancer cells with a *bottom-of-the-crypt* phenotype (EpCAM^+^, CD44^+^, CD166^+^) purified by FACS from the PDX-COLON-8 line following infection with either an empty lentivirus vector or lentivirus vectors encoding for the E2F4-shRNA [#1] or E2F4-shRNA [#2] constructs. The analysis confirmed the capacity of both shRNAs to cause a visually detectable reduction in E2F4 protein levels in the population with a *bottom-of-the-crypt* phenotype (EpCAM^+^, CD44^+^, CD166^+^), which is enriched in cells with *cancer stem cell* (CSC) properties. ACTB (β-Actin) protein levels were analyzed in parallel and used as a visual standard to normalize for the total amount of protein loaded onto the gel.

**Supplementary Figure S4.**
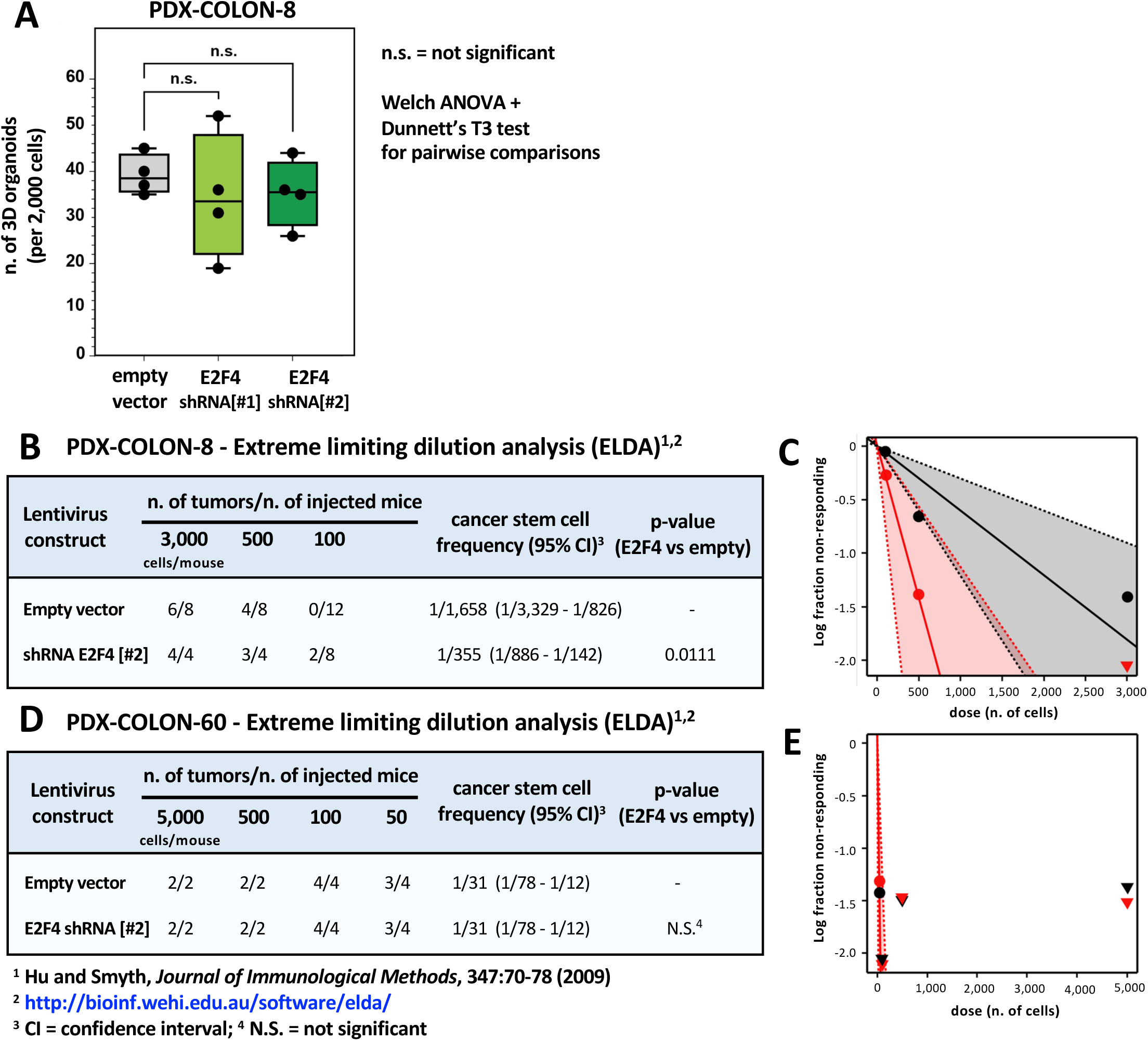
Downregulation of E2F4 expression does not result in reductions of either the *in vitro* organoid-forming capacity or the *in vivo* tumor-forming capacity of human colon cancer cells. The role of E2F4 in regulating the tumor-initiating capacity of colon cancer cells was evaluated in two *patient-derived xenograft* (PDX) lines infected with lentivirus vectors encoding E2F4-specific *short-hairpin RNA* (shRNA) constructs in tandem with a fluorescent reporter (copGFP). Infected cancer cells were sorted based on copGFP expression and tested prospectively for changes in their capacity to form 3D organoids (*in vitro*) or solid tumors (*in vivo*) as compared to cells infected with an empty vector. **(A)** Comparison of the number of 3D organoids formed by PDX-COLON-8 cancer cells infected with different lentivirus vectors, after plating on Matrigel (n=2,000 cells/well). n.s.: not significant (Welch ANOVA + Dunnett’s T3 test for pairwise comparisons). (B-E) *Extreme limiting dilution analysis* (ELDA) of the *in vivo* tumorigenic capacity of cancer cells from either the PDX-COLON-8 (B-C) or the PDX-COLON-60 (D-E) lines, following infection with lentivirus vectors encoding E2F4-shRNA constructs. The ELDA was based on the approach described by Hu & Smyth (*J. Immunol. Methods*, 347:70-78, 2009). Cancer cells infected with E2F4-shRNA constructs did not display reductions in their capacity to form organoids (*in vitro*) or solid tumors (*in vivo*) as compared to cells infected with an empty vector.

**Supplementary Table 1:**
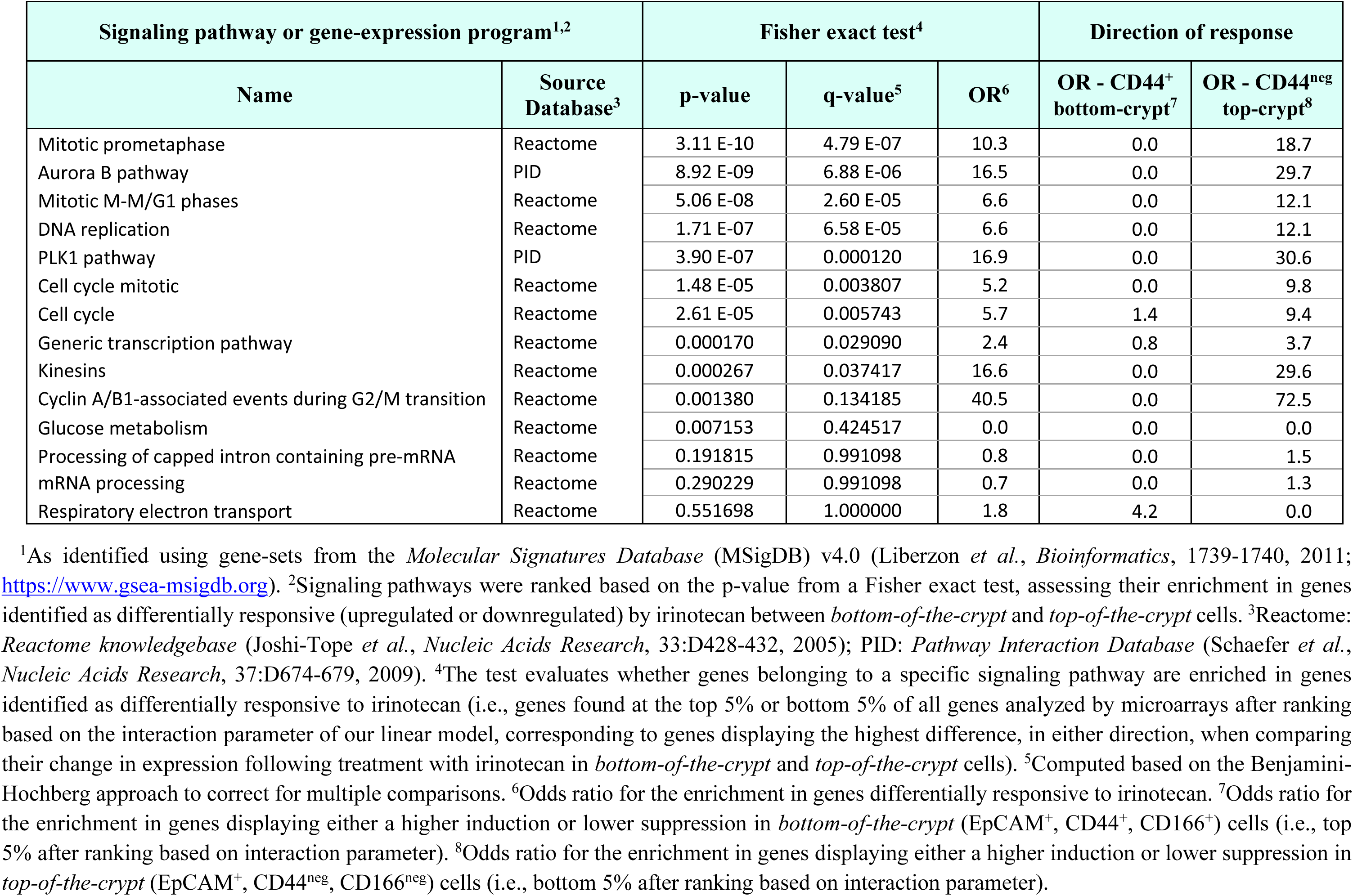
Signaling pathways and gene-expression programs identified as differentially modulated between colon cancer cells with a *bottom-of-the-crypt* (EpCAM^+^, CD44^+^, CD166^+^) and a *top-of-the-crypt* (EpCAM^+^, CD44^neg^, CD166^neg^) phenotype, following *in vivo* treatment with irinotecan (CPT-11).

| Signaling pathway or gene-expression program<sup>1,2</sup> |  | Fisher exact test<sup>4</sup> |  |  | Direction of response |  |
| --- | --- | --- | --- | --- | --- | --- |
| Name | Source Database<sup>3</sup> | p-value | q-value<sup>5</sup> | OR<sup>6</sup> | OR - CD44<sup>+</sup> bottom-crypt<sup>7</sup> | OR - CD44<sup>neg</sup> top-crypt<sup>8</sup> |
| Mitotic prometaphase | Reactome | 3.11 E-10 | 4.79 E-07 | 10.3 | 0.0 | 18.7 |
| Aurora B pathway | PID | 8.92 E-09 | 6.88 E-06 | 16.5 | 0.0 | 29.7 |
| Mitotic M-M/G1 phases | Reactome | 5.06 E-08 | 2.60 E-05 | 6.6 | 0.0 | 12.1 |
| DNA replication | Reactome | 1.71 E-07 | 6.58 E-05 | 6.6 | 0.0 | 12.1 |
| PLK1 pathway | PID | 3.90 E-07 | 0.000120 | 16.9 | 0.0 | 30.6 |
| Cell cycle mitotic | Reactome | 1.48 E-05 | 0.003807 | 5.2 | 0.0 | 9.8 |
| Cell cycle | Reactome | 2.61 E-05 | 0.005743 | 5.7 | 1.4 | 9.4 |
| Generic transcription pathway | Reactome | 0.000170 | 0.029090 | 2.4 | 0.8 | 3.7 |
| Kinesins | Reactome | 0.000267 | 0.037417 | 16.6 | 0.0 | 29.6 |
| Cyclin A/B1-associated events during G2/M transition | Reactome | 0.001380 | 0.134185 | 40.5 | 0.0 | 72.5 |
| Glucose metabolism | Reactome | 0.007153 | 0.424517 | 0.0 | 0.0 | 0.0 |
| Processing of capped intron containing pre-mRNA | Reactome | 0.191815 | 0.991098 | 0.8 | 0.0 | 1.5 |
| mRNA processing | Reactome | 0.290229 | 0.991098 | 0.7 | 0.0 | 1.3 |
| Respiratory electron transport | Reactome | 0.551698 | 1.000000 | 1.8 | 4.2 | 0.0 |

**Supplementary Table 3:** Clinical and pathological characteristics of the human *colorectal carcinomas* (CRCs) from which the *patient derived xenograft* (PDX) models utilized in this study were established.

| PDX line | Patient |  | primary site of origin | primary vs. metastasis | Tumor stage<sup>1</sup> |  | Tumor grade<sup>2</sup> |
| --- | --- | --- | --- | --- | --- | --- | --- |
|  | age | sex |  |  | TNM | AJCC |  |
| PDX-COLON-8 | 49 | male | Sigmoid colon | primary | T3N0 | IIa | G2 |
| PDX-COLON-60 | 58 | male | Right colon | primary | T4aN2bM1b | IVb | G2 |
| PDX-COLON-136 | 74 | female | Right colon | primary | T4N1bM0 | IIIb | G3 |
| PDX-COLON-441 | n.a. | male | Left colon | primary | T3N2bM0 | IIIc | n.a. |

<sup>1</sup> *Tumor-Node-Metastasis* (TNM) status and pathological stage are reported according to the 7<sup>th</sup> edition of the *American Joint Committee on Cancer* (AJCC) staging manual for colorectal cancer (2009). <sup>2</sup> Grade of histological differentiation, annotated as follows: G1, well differentiated; G2: moderately differentiated; G3, poorly differentiated; n.a., not available.

**Supplementary Table 4:**
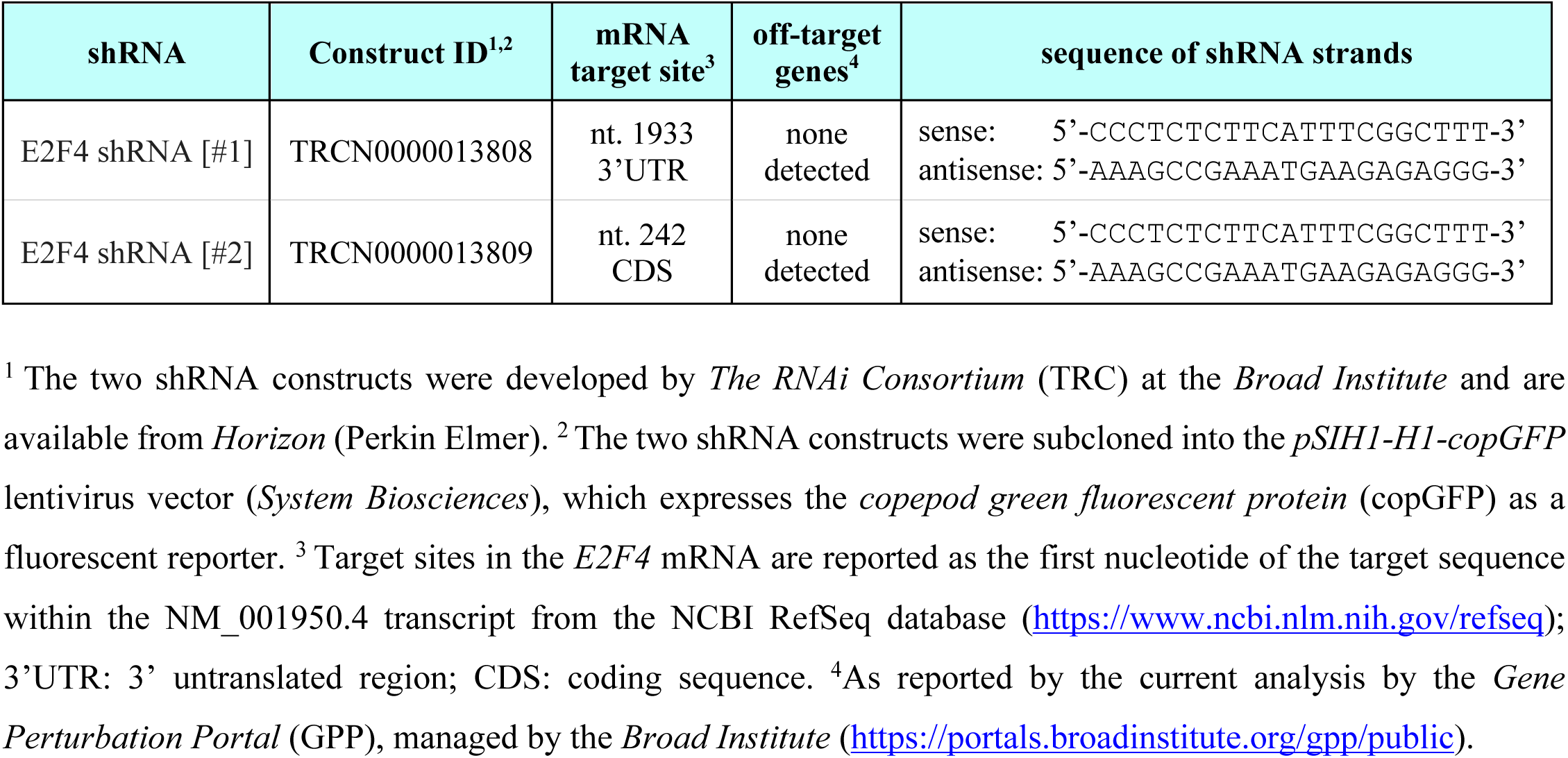
Molecular properties of the *short hairpin RNA* (shRNA) constructs used to knock-down E2F4 protein expression in this study.

## REFERENCES

1. Bray F, Laversanne M, Sung H, Ferlay J, Siegel RL, Soerjomataram I, Jemal A: Global cancer statistics 2022: GLOBOCAN estimates of incidence and mortality worldwide for 36 cancers in 185 countries. CA Cancer Journal for Clinicians, 74:229-263 (2024)

2. van Gestel YR, de Hingh IH, van Herk-Sukel MP, van Erning FN, Beerepoot LV, Wijsman JH, Slooter GD, Rutten HJ, Creemers GJ, Lemmens VE: Patterns of metachronous metastases after curative treatment of colorectal cancer. Cancer Epidemiology, 38:448–454 (2014)

3. Vayrynen V, Wirta EV, Seppala T, Sihvo E, Mecklin JP, Vasala K, Kellokumpu I: Incidence and management of patients with colorectal cancer and synchronous and metachronous colorectal metastases: a population-based study. BJS Open, 4:685–692 (2020)

4. Morris VK, Kennedy EB, Baxter NN, Benson AB, 3rd, Cercek A, Cho M, Ciombor KK, Cremolini C, Davis A, Deming DA et al: Treatment of Metastatic Colorectal Cancer: ASCO Guideline. Journal of Clinical Oncology, 41:678–700 (2023)

5. Cervantes A, Adam R, Rosello S, Arnold D, Normanno N, Taieb J, Seligmann J, De Baere T, Osterlund P, Yoshino T et al: Metastatic colorectal cancer: ESMO Clinical Practice Guideline for diagnosis, treatment and follow-up. Annals of Oncology, 34:10–32 (2023)

6. O’Connell JB, Maggard MA, Ko CY: Colon cancer survival rates with the new American Joint Committee on Cancer sixth edition staging. Journal of the National Cancer Institute (J.N.C.I.), 96:1420–1425 (2004)

7. Neugut AI, Lin A, Raab GT, Hillyer GC, Keller D, O’Neil DS, Accordino MK, Kiran RP, Wright J, Hershman DL: FOLFOX and FOLFIRI Use in Stage IV Colon Cancer: Analysis of SEER-Medicare Data. Clinical Colorectal Cancer, 18:133–140 (2019)

8. Dalerba P, Dylla SJ, Park IK, Liu R, Wang X, Cho RW, Hoey T, Gurney A, Huang EH, Simeone DM et al: Phenotypic characterization of human colorectal cancer stem cells. Proceedings of the National Academy of Sciences USA (P.N.A.S.), 104:10158–10163 (2007)

9. Dalerba P, Kalisky T, Sahoo D, Rajendran PS, Rothenberg ME, Leyrat AA, Sim S, Okamoto J, Johnston DM, Qian D et al: Single-cell dissection of transcriptional heterogeneity in human colon tumors. Nature Biotechnology, 29:1120–1127 (2011)

10. Dalerba P, Cho RW, Clarke MF: Cancer stem cells: models and concepts. Annual Review of Medicine, 58:267–284 (2007)

11. Ricci-Vitiani L, Lombardi DG, Pilozzi E, Biffoni M, Todaro M, Peschle C, De Maria R: Identification and expansion of human colon-cancer-initiating cells. Nature, 445:111–115 (2007)

12. Dylla SJ, Beviglia L, Park IK, Chartier C, Raval J, Ngan L, Pickell K, Aguilar J, Lazetic S, Smith-Berdan S et al: Colorectal cancer stem cells are enriched in xenogeneic tumors following chemotherapy. PLoS One, 3:e2428 (2008)

13. Chen J, Li Y, Yu TS, McKay RM, Burns DK, Kernie SG, Parada LF: A restricted cell population propagates glioblastoma growth after chemotherapy. Nature, 488:522–526 (2012)

14. Bao S, Wu Q, McLendon RE, Hao Y, Shi Q, Hjelmeland AB, Dewhirst MW, Bigner DD, Rich JN: Glioma stem cells promote radioresistance by preferential activation of the DNA damage response. Nature, 444:756–760 (2006)

15. Diehn M, Cho RW, Lobo NA, Kalisky T, Dorie MJ, Kulp AN, Qian D, Lam JS, Ailles LE, Wong M et al: Association of reactive oxygen species levels and radioresistance in cancer stem cells. Nature, 458:780–783 (2009)

16. Cunningham D, Humblet Y, Siena S, Khayat D, Bleiberg H, Santoro A, Bets D, Mueser M, Harstrick A, Verslype C et al: Cetuximab monotherapy and cetuximab plus irinotecan in irinotecan-refractory metastatic colorectal cancer. The New England Journal of Medicine (N.E.J.M.), 351:337–345 (2004)

17. Saltz LB, Cox JV, Blanke C, Rosen LS, Fehrenbacher L, Moore MJ, Maroun JA, Ackland SP, Locker PK, Pirotta N et al: Irinotecan plus fluorouracil and leucovorin for metastatic colorectal cancer. Irinotecan Study Group. The New England Journal of Medicine (N.E.J.M.), 343:905–914 (2000)

18. Li YF, Altman RB: Systematic target function annotation of human transcription factors. BMC Biology, 16:4 (2018)

19. Burridge PW, Li YF, Matsa E, Wu H, Ong SG, Sharma A, Holmstrom A, Chang AC, Coronado MJ, Ebert AD et al: Human induced pluripotent stem cell-derived cardiomyocytes recapitulate the predilection of breast cancer patients to doxorubicin-induced cardiotoxicity. Nature Medicine, 22:547–556 (2016)

20. Nguyen PK, Lee WH, Li YF, Hong WX, Hu S, Chan C, Liang G, Nguyen I, Ong SG, Churko J et al: Assessment of the Radiation Effects of Cardiac CT Angiography Using Protein and Genetic Biomarkers. JACC Cardiovascular Imaging, 8:873–884 (2015)

21. Rothenberg ME, Nusse Y, Kalisky T, Lee JJ, Dalerba P, Scheeren F, Lobo N, Kulkarni S, Sim S, Qian D et al: Identification of a cKit(+) colonic crypt base secretory cell that supports Lgr5(+) stem cells in mice. Gastroenterology, 142:1195–1205 (2012)

22. Barker N, van Es JH, Kuipers J, Kujala P, van den Born M, Cozijnsen M, Haegebarth A, Korving J, Begthel H, Peters PJ et al: Identification of stem cells in small intestine and colon by marker gene Lgr5. Nature, 449:1003–1007 (2007)

23. Capdevila C, Miller J, Cheng L, Kornberg A, George JJ, Lee H, Botella T, Moon CS, Murray JW, Lam S et al: Time-resolved fate mapping identifies the intestinal upper crypt zone as an origin of Lgr5+ crypt base columnar cells. Cell, 187:3039–3055 (2024)

24. Malagola E, Vasciaveo A, Ochiai Y, Kim W, Zheng B, Zanella L, Wang ALE, Middelhoff M, Nienhuser H, Deng L et al: Isthmus progenitor cells contribute to homeostatic cellular turnover and support regeneration following intestinal injury. Cell, 187:3056–3071 (2024)

25. Moffat J, Grueneberg DA, Yang X, Kim SY, Kloepfer AM, Hinkle G, Piqani B, Eisenhaure TM, Luo B, Grenier JK et al: A lentiviral RNAi library for human and mouse genes applied to an arrayed viral high-content screen. Cell, 124:1283–1298 (2006)

26. Conboy CM, Spyrou C, Thorne NP, Wade EJ, Barbosa-Morais NL, Wilson MD, Bhattacharjee A, Young RA, Tavare S, Lees JA et al: Cell cycle genes are the evolutionarily conserved targets of the E2F4 transcription factor. PLoS One, 2:e1061 (2007)

27. Lee BK, Bhinge AA, Iyer VR: Wide-ranging functions of E2F4 in transcriptional activation and repression revealed by genome-wide analysis. Nucleic Acids Research, 39:3558–3573 (2011)

28. Sadasivam S, DeCaprio JA: The DREAM complex: master coordinator of cell cycle-dependent gene expression. Nature Reviews Cancer, 13:585–595 (2013)

29. Plesca D, Crosby ME, Gupta D, Almasan A: E2F4 function in G2: maintaining G2-arrest to prevent mitotic entry with damaged DNA. Cell Cycle, 6:1147–1152 (2007)

30. Crosby ME, Jacobberger J, Gupta D, Macklis RM, Almasan A: E2F4 regulates a stable G2 arrest response to genotoxic stress in prostate carcinoma. Oncogene, 26:1897–1909 (2007)

31. Wang P, Karakose E, Argmann C, Wang H, Balev M, Brody RI, Rivas HG, Liu X, Wood O, Liu H et al: Disrupting the DREAM complex enables proliferation of adult human pancreatic beta cells. Journal of Clinical Investigation, 132:e157086 (2022)

32. Mathijssen RH, van Alphen RJ, Verweij J, Loos WJ, Nooter K, Stoter G, Sparreboom A: Clinical pharmacokinetics and metabolism of irinotecan (CPT-11). Clinical Cancer Research, 7:2182–2194 (2001)

33. Hsiang YH, Liu LF, Wall ME, Wani MC, Nicholas AW, Manikumar G, Kirschenbaum S, Silber R, Potmesil M: DNA topoisomerase I-mediated DNA cleavage and cytotoxicity of camptothecin analogues. Cancer Research, 49:4385–4389 (1989)

34. Kawato Y, Aonuma M, Hirota Y, Kuga H, Sato K: Intracellular roles of SN-38, a metabolite of the camptothecin derivative CPT-11, in the antitumor effect of CPT-11. Cancer Research, 51:4187–4191 (1991)

35. Pommier Y, Nussenzweig A, Takeda S, Austin C: Human topoisomerases and their roles in genome stability and organization. Nature Reviews Molecular & Cell Biology, 23:407–427 (2022)

36. Liu LF, Desai SD, Li TK, Mao Y, Sun M, Sim SP: Mechanism of action of camptothecin. Annals of the New York Academy of Sciences, 922:1–10 (2000)

37. Liu LF, Duann P, Lin CT, D’Arpa P, Wu J: Mechanism of action of camptothecin. Annals of the New York Academy of Sciences, 803:44–49 (1996)

38. Covey JM, Jaxel C, Kohn KW, Pommier Y: Protein-linked DNA strand breaks induced in mammalian cells by camptothecin, an inhibitor of topoisomerase I. Cancer Research, 49:5016–5022 (1989)

39. Mattern MR, Mong SM, Bartus HF, Mirabelli CK, Crooke ST, Johnson RK: Relationship between the intracellular effects of camptothecin and the inhibition of DNA topoisomerase I in cultured L1210 cells. Cancer Research, 47:1793–1798 (1987)

40. Holm C, Covey JM, Kerrigan D, Pommier Y: Differential requirement of DNA replication for the cytotoxicity of DNA topoisomerase I and II inhibitors in Chinese hamster DC3F cells. Cancer Research, 49:6365–6368 (1989)

41. D’Arpa P, Beardmore C, Liu LF: Involvement of nucleic acid synthesis in cell killing mechanisms of topoisomerase poisons. Cancer Research, 50:6919–6924 (1990)

42. Deyme L, Barbolosi D, Mbatchi LC, Tubiana-Mathieu N, Ychou M, Evrard A, Gattacceca F: Population pharmacokinetic model of irinotecan and its four main metabolites in patients treated with FOLFIRI or FOLFIRINOX regimen. Cancer Chemotherapy and Pharmacology, 88:247–258 (2021)

43. Tanizawa A, Fujimori A, Fujimori Y, Pommier Y: Comparison of topoisomerase I inhibition, DNA damage, and cytotoxicity of camptothecin derivatives presently in clinical trials. Journal of the National Cancer Institute (J.N.C.I.), 86:836–842 (1994)

44. Rempel RE, Saenz-Robles MT, Storms R, Morham S, Ishida S, Engel A, Jakoi L, Melhem MF, Pipas JM, Smith C et al: Loss of E2F4 activity leads to abnormal development of multiple cellular lineages. Molecular Cell, 6:293–306 (2000)

45. Paquin MC, Leblanc C, Lemieux E, Bian B, Rivard N: Functional impact of colorectal cancer-associated mutations in the transcription factor E2F4. International Journal of Oncology, 43:2015–2022 (2013)

46. Laham AJ, El-Awady R, Saber-Ayad M, Wang N, Yan G, Boudreault J, Ali S, Lebrun JJ: Targeting the DYRK1A kinase prevents cancer progression and metastasis and promotes cancer cells response to G1/S targeting chemotherapy drugs. NPJ Precision Oncology, 8:128 (2024)

47. Wang P, Alvarez-Perez JC, Felsenfeld DP, Liu H, Sivendran S, Bender A, Kumar A, Sanchez R, Scott DK, Garcia-Ocana A et al: A high-throughput chemical screen reveals that harmine-mediated inhibition of DYRK1A increases human pancreatic beta cell replication. Nature Medicine, 21:383–388 (2015)

48. Ables JL, Israel L, Wood O, Govindarajulu U, Fremont RT, Banerjee R, Liu H, Cohen J, Wang P, Kumar K et al: A Phase 1 single ascending dose study of pure oral harmine in healthy volunteers. Journal of Psychopharmacology, 38:911–923 (2024)

49. Callaway JC, McKenna DJ, Grob CS, Brito GS, Raymon LP, Poland RE, Andrade EN, Andrade EO, Mash DC: Pharmacokinetics of Hoasca alkaloids in healthy humans. Journal of Ethnopharmacology, 65:243–256 (1999)

50. ten Bokkel Huinink W, Lane SR, Ross GA, International Topotecan Study G: Long-term survival in a phase III, randomised study of topotecan versus paclitaxel in advanced epithelial ovarian carcinoma. Annals of Oncology, 15:100-103 (2004)

51. Adams E, Wildiers H, Neven P, Punie K: Sacituzumab govitecan and trastuzumab deruxtecan: two new antibody-drug conjugates in the breast cancer treatment landscape. ESMO Open, 6:100204 (2021)

52. Aston WJ, Hope DE, Nowak AK, Robinson BW, Lake RA, Lesterhuis WJ: A systematic investigation of the maximum tolerated dose of cytotoxic chemotherapy with and without supportive care in mice. BMC Cancer, 17:684 (2017)

53. Bissery MC, Vrignaud P, Lavelle F, Chabot GG: Experimental antitumor activity and pharmacokinetics of the camptothecin analog irinotecan (CPT-11) in mice. Anticancer Drugs, 7:437–460 (1996)

54. Guichard S, Chatelut E, Lochon I, Bugat R, Mahjoubi M, Canal P: Comparison of the pharmacokinetics and efficacy of irinotecan after administration by the intravenous versus intraperitoneal route in mice. Cancer Chemotherapy and Pharmacology, 42:165–170 (1998)

55. Tomayko MM, Reynolds CP: Determination of subcutaneous tumor size in athymic (nude) mice. Cancer Chemotherapy and Pharmacology, 24:148–154 (1989)

56. Hather G, Liu R, Bandi S, Mettetal J, Manfredi M, Shyu WC, Donelan J, Chakravarty A: Growth rate analysis and efficient experimental design for tumor xenograft studies. Cancer Informatics, 13(Suppl 4):65–72 (2014)

57. Oberg AL, Heinzen EP, Hou X, Al Hilli MM, Hurley RM, Wahner Hendrickson AE, Goergen KM, Larson MC, Becker MA, Eckel-Passow JE et al: Statistical analysis of comparative tumor growth repeated measures experiments in the ovarian cancer patient derived xenograft (PDX) setting. Scientific Reports, 11:8076 (2021)

58. Viragova S, Aparicio L, Palmerini P, Zhao J, Valencia Salazar LE, Schurer A, Dhuri A, Sahoo D, Moskaluk CA, Rabadan R et al: Inverse agonists of retinoic acid receptor/retinoid X receptor signaling as lineage-specific antitumor agents against human adenoid cystic carcinoma. Journal of the National Cancer Institute (J.N.C.I.), 115:838–852 (2023)

59. Hisamori S, Mukohyama J, Koul S, Hayashi T, Rothenberg ME, Maeda M, Isobe T, Valencia Salazar LE, Qian X, Johnston DM et al: Upregulation of BMI1-suppressor miRNAs (miR-200c, miR-203) during terminal differentiation of colon epithelial cells. Journal of Gastroenterology, 57:407–422 (2022)

60. Dunnett CW: Pairwise Multiple Comparisons in the Unequal Variance Case. Journal of the American Statistical Association, 75:796-800 (1980)

61. Irizarry RA, Hobbs B, Collin F, Beazer-Barclay YD, Antonellis KJ, Scherf U, Speed TP: Exploration, normalization, and summaries of high density oligonucleotide array probe level data. Biostatistics (Oxford, England), 4:249–264 (2003)

62. Bolstad BM, Irizarry RA, Astrand M, Speed TP: A comparison of normalization methods for high density oligonucleotide array data based on variance and bias. Bioinformatics, 19:185–193 (2003)

63. Jiang Z, Gentleman R: Extensions to gene set enrichment. Bioinformatics, 23:306–313 (2007)

64. Oron AP, Jiang Z, Gentleman R: Gene set enrichment analysis using linear models and diagnostics. Bioinformatics, 24:2586–2591 (2008)

65. Ackermann M, Strimmer K: A general modular framework for gene set enrichment analysis. BMC Bioinformatics, 10:47 (2009)

66. Liberzon A, Subramanian A, Pinchback R, Thorvaldsdottir H, Tamayo P, Mesirov JP: Molecular signatures database (MSigDB) 3.0. Bioinformatics, 27:1739-1740 (2011)

67. Joshi-Tope G, Gillespie M, Vastrik I, D’Eustachio P, Schmidt E, de Bono B, Jassal B, Gopinath GR, Wu GR, Matthews L et al: Reactome: a knowledgebase of biological pathways. Nucleic Acids Research, 33(Database):D428-432 (2005)

68. Schaefer CF, Anthony K, Krupa S, Buchoff J, Day M, Hannay T, Buetow KH: PID: the Pathway Interaction Database. Nucleic Acids Research, 37(Database):D674-679 (2009)

69. Li YF, Xin F, Altman RB: Separating the Causes and Consequences in Disease Transcriptome. Pacific Symposium on Biocomputing, 21:381–392 (2016)

70. Benjamini Y, Hochberg Y: Controlling the False Discovery Rate: A Practical and Powerful Approach to Multiple Testing. Journal of the Royal Statistical Society Series B: Statistical Methodology, 57:289-300 (1995)

71. Cui X, Churchill GA: Statistical tests for differential expression in cDNA microarray experiments. Genome Biology, 4:210 (2003)

72. Cancer Genome Atlas N: Comprehensive molecular characterization of human colon and rectal cancer. Nature, 487:330-337 (2012)

73. Liu J, Lichtenberg T, Hoadley KA, Poisson LM, Lazar AJ, Cherniack AD, Kovatich AJ, Benz CC, Levine DA, Lee AV et al: An Integrated TCGA Pan-Cancer Clinical Data Resource to Drive High-Quality Survival Outcome Analytics. Cell, 173:400–416 (2018)

74. Hintze JL, Nelson RD: Violin Plots: A Box Plot-Density Trace Synergism. The American Statistician, 52:181-184 (1998)

75. Williamson DF, Parker RA, Kendrick JS: The box plot: a simple visual method to interpret data. Annals of Internal Medicine, 110:916–921 (1989)

76. Schroeder A, Mueller O, Stocker S, Salowsky R, Leiber M, Gassmann M, Lightfoot S, Menzel W, Granzow M, Ragg T: The RIN: an RNA integrity number for assigning integrity values to RNA measurements. BMC Molecular Biology, 7:3 (2006)

77. Fleige S, Pfaffl MW: RNA integrity and the effect on the real-time qRT-PCR performance. Molecular Aspects of Medicine, 27:126–139 (2006)

78. Wu J, Anczukow O, Krainer AR, Zhang MQ, Zhang C: OLego: fast and sensitive mapping of spliced mRNA-Seq reads using small seeds. Nucleic Acids Research, 41:5149–5163 (2013)

79. Trapnell C, Pachter L, Salzberg SL: TopHat: discovering splice junctions with RNA-Seq. Bioinformatics, 25:1105–1111 (2009)

80. Trapnell C, Roberts A, Goff L, Pertea G, Kim D, Kelley DR, Pimentel H, Salzberg SL, Rinn JL, Pachter L: Differential gene and transcript expression analysis of RNA-seq experiments with TopHat and Cufflinks. Nature Protocols, 7:562–578 (2012)

81. Brattain MG, Fine WD, Khaled FM, Thompson J, Brattain DE: Heterogeneity of malignant cells from a human colonic carcinoma. Cancer Research, 41:1751–1756 (1981)

82. von Kleist S, Chany E, Burtin P, King M, Fogh J: Immunohistology of the antigenic pattern of a continuous cell line from a human colon tumor. Journal of the National Cancer Institute (J.N.C.I.), 55:555–560 (1975)

83. Kawai K, Viars C, Arden K, Tarin D, Urquidi V, Goodison S: Comprehensive karyotyping of the HT-29 colon adenocarcinoma cell line. Genes Chromosomes Cancer, 34:1–8 (2002)

84. Tiscornia G, Singer O, Verma IM: Production and purification of lentiviral vectors. Nature Protocols, 1:241–245 (2006)

85. Ailles LE, Naldini L: HIV-1-derived lentiviral vectors. Current Topics in Microbiology Immunology, 261:31–52 (2002)

86. Dull T, Zufferey R, Kelly M, Mandel RJ, Nguyen M, Trono D, Naldini L: A third-generation lentivirus vector with a conditional packaging system. Journal of Virology, 72:8463–8471 (1998)

87. Zufferey R, Dull T, Mandel RJ, Bukovsky A, Quiroz D, Naldini L, Trono D: Self-inactivating lentivirus vector for safe and efficient in vivo gene delivery. Journal of Virology, 72:9873–9880 (1998)

88. Miyoshi H, Blomer U, Takahashi M, Gage FH, Verma IM: Development of a self-inactivating lentivirus vector. Journal of Virology, 72:8150–8157 (1998)

89. Scherr M, Battmer K, Ganser A, Eder M: Modulation of gene expression by lentiviral-mediated delivery of small interfering RNA. Cell Cycle, 2:251–257 (2003)

90. Kingston RE, Chen CA, Okayama H: Calcium phosphate transfection. Current Protocols in Cell Biology, Chapter 20, Unit 20.3 (2003)

91. Ellis BL, Potts PR, Porteus MH: Creating higher titer lentivirus with caffeine. Human Gene Therapy, 22:93–100 (2011)

92. O’Doherty U, Swiggard WJ, Malim MH: Human immunodeficiency virus type 1 spinoculation enhances infection through virus binding. Journal of Virology, 74:10074–10080 (2000)

93. Bondanza A, Valtolina V, Magnani Z, Ponzoni M, Fleischhauer K, Bonyhadi M, Traversari C, Sanvito F, Toma S, Radrizzani M et al: Suicide gene therapy of graft-versus-host disease induced by central memory human T lymphocytes. Blood, 107:1828–1836 (2006)

94. Sato T, Stange DE, Ferrante M, Vries RG, Van Es JH, Van den Brink S, Van Houdt WJ, Pronk A, Van Gorp J, Siersema PD et al: Long-term expansion of epithelial organoids from human colon, adenoma, adenocarcinoma, and Barrett’s epithelium. Gastroenterology, 141:1762–1772 (2011)

95. Shimono Y, Mukohyama J, Isobe T, Johnston DM, Dalerba P, Suzuki A: Organoid Culture of Human Cancer Stem Cells. Methods in Molecular Biology, 1576:23–31 (2019)

96. Livak KJ, Schmittgen TD: Analysis of relative gene expression data using real-time quantitative PCR and the 2-^Delta Delta CT^ Method. Methods, 25:402–408 (2001)

97. Chomczynski P, Sacchi N: Single-step method of RNA isolation by acid guanidinium thiocyanate-phenol-chloroform extraction. Analytical Biochemistry, 162:156–159 (1987)

98. Chomczynski P, Sacchi N: The single-step method of RNA isolation by acid guanidinium thiocyanate-phenol-chloroform extraction: twenty-something years on. Nature Protocols, 1:581–585 (2006)

99. Honda K, Yamada T, Hayashida Y, Idogawa M, Sato S, Hasegawa F, Ino Y, Ono M, Hirohashi S: Actinin-4 increases cell motility and promotes lymph node metastasis of colorectal cancer. Gastroenterology, 128:51–62 (2005)

100. Matsubara J, Ono M, Negishi A, Ueno H, Okusaka T, Furuse J, Furuta K, Sugiyama E, Saito Y, Kaniwa N et al: Identification of a predictive biomarker for hematologic toxicities of gemcitabine. Journal of Clinical Oncology, 27:2261–2268 (2009)

101. Hu Y, Smyth GK: ELDA: extreme limiting dilution analysis for comparing depleted and enriched populations in stem cell and other assays. Journal of Immunological Methods, 347:70-78 (2009)

102. Kusser KL, Randall TD: Simultaneous detection of EGFP and cell surface markers by fluorescence microscopy in lymphoid tissues. Journal of Histochemistry and Cytochemistry, 51:5–14 (2003)

